# Mi-2 and E93 coordinate robust cell cycle exit with terminal differentiation through enhancer decommissioning

**DOI:** 10.64898/2026.09.16.752104

**Authors:** Elli M. Buchert, Elizabeth Fogarty, Yiqin Ma, Saurabh Jayesh Kumar Mehta, Daniel J. McKay, Laura A. Buttitta

## Abstract

Many postmitotic tissues coordinate robust cell cycle exit with the progression of terminal differentiation. While the signals involved in initiating cell cycle exit during terminal differentiation for some tissues have been described, less is known about the mechanisms that maintain a stable, non-cycling state. We previously found that a subset of rate-limiting cell cycle genes are subject to developmentally controlled changes in chromatin accessibility after cell cycle exit, suggesting chromatin remodelers may play a key role in maintaining a robust postmitotic state during terminal differentiation. Here we show that the general chromatin remodeler Mi-2 is required to ensure a stable postmitotic state in *Drosophila* eyes and wings. Mi-2-mediated remodeling coordinates cell state transitions during development, as compromising Mi-2 function slows the chromatin accessibility and gene expression dynamics that are critical for tissue maturation. Mi-2 and a temporally expressed transcription factor involved in terminal differentiation, E93, each decommission chromatin at developmentally dynamic regions near a limited set of shared target genes, including the early differentiation factor *broad* and the rate-limiting mitotic regulator *string* (*cdc25c*). Our evidence supports a model where Mi-2 and E93 act *in vivo* to decommission a newly identified distal *string* enhancer that is active during cell cycle exit, to coordinate the transition to a robust postmitotic state with the progression of terminal differentiation. Temporally regulated enhancer decommissioning at the *string* locus, depending on both an instructive temporal factor and an enzymatic remodeler, provides a molecular explanation for the long-term, nearly irreversible postmitotic state observed in many tissues as terminal differentiation progresses.

## Introduction

*Drosophila melanogaster* metamorphosis is a time of dramatic developmental changes, during which many larval tissues are completely remodeled to allow for morphogenesis of the adult structures. The imaginal discs giving rise to the adult eyes, wings, and legs undergo temporally coordinated morphological and cell cycle changes during this time. The temporal coordination occurs in response to a pulse of the steroid hormone ecdysone, which triggers the larval to puparium transition, an event marking the beginning of metamorphosis [1]. This transition sets in place a “feed-forward” cascade of ecdysone-induced temporal transcriptional changes that coordinate events across tissues. This feed-forward system works by sequential activation of early-responsive targets that induce later targets, concurrent with negative feedback inhibition of the early response, resulting in a unidirectional transcriptional cascade that robustly drives developmental transitions [2]. This model serves as a prototype for many examples across organisms, where temporally transient signals generate sequential and robustly irreversible cell state changes.

The ecdysone transcriptional cascade has been best described for the salivary gland, where the large polytene chromosomes provide a visible, temporal readout of target activation through chromosome “puffs” at the genomic locations of ecdysone target genes [3]. However, it is becoming clear that the order and identity of the genes in the ecdysone transcriptional response are tissue specific, which is likely essential for shaping tissue-specific responses to the systemic pulses of ecdysone [4–6]. For example, in abdominal histoblasts, cell proliferation is induced during early metamorphosis in response to ecdysone through ecdysone-dependent inhibition of a microRNA suppressor of the mitotic regulator Cdc25c, known as String in flies [7, 8]. By contrast, in the wing the same ecdysone pulse induces the transcriptional repressor Broad which temporarily suppresses expression of *string* leading to a transient G2 arrest that synchronizes the cell cycle across the tissue during metamorphosis [4]. Thus, the transcriptional response to the hormone is tailored in a tissue-specific manner, such that the same systemic pulse of ecdysone can simultaneously induce divergent outcomes in different parts of the organism.

A second, larger pulse of ecdysone begins around 24 hours into metamorphosis. Coincident with this, the imaginal tissues giving rise to the adult eyes, wings, and legs undergo their final cell cycles and a temporally coordinated, permanent cell cycle exit [9–12]. The role for ecdysone in promoting entry into a permanent postmitotic state at this stage of development is unclear, although ecdysone targets that are activated at this time such as the E93 transcription factor are known to be involved in tissue maturation and terminal differentiation, suggesting the possibility of coordination between the two processes [6].

We previously performed chromatin accessibility and transcriptomic analyses of pupal eyes and wings during cell cycle arrest and terminal differentiation with a goal to identify gene regulatory networks coordinating the two processes [13, 14]. We found that most cell cycle genes (such as *pcna* and *Orc6*) maintain accessibility at their promoters and promoter proximal regulatory elements despite transitioning to a transcriptionally repressed after cell cycle exit. By contrast, three critical rate-limiting cell cycle regulatory genes with complex, modular enhancers spanning many kilobases exhibit regulatory element closing after cell cycle exit, potentially making them refractory to reactivation after cell cycle exit [13]. The mechanisms by which these key cis-regulatory elements close during cell cycle exit remain unclear. Here, we investigate whether chromatin remodelers play roles in promoting and maintaining robust cell cycle exit during metamorphosis via control of chromatin accessibility. Through a candidate genetic screen, we identified a role for the chromatin remodeler Mi-2 in promoting cell cycle exit during metamorphosis. Mi-2 is a homolog of the mammalian CHD3/4 proteins, which are ATP-dependent chromatin remodelers of the Chromodomain-Helicase-DNA (CHD) binding family [15]. Other ATP-dependent chromatin remodelers such as SWI/SNF have been shown to function in opening chromatin at differentiation genes to allow for gene activation but have also been shown to regulate cell cycle exit in other organisms by contributing to the repression of positive cell cycle regulators [16–18].

We further sought to determine how the Mi-2 chromatin remodeler affects cell cycle exit by examining chromatin accessibility and gene expression changes when Mi-2 function is compromised. We show that Mi-2 supports timely developmental dynamics including chromatin accessibility and gene expression changes as cells exit the cell cycle and proceed with terminal differentiation. Our data suggest that Mi-2 and the temporal factor E93 may coordinate dynamics at a limited overlapping set of cell cycle and differentiation genes to ensure coordination between postmitotic cell cycle exit and terminal differentiation. We identify a previously uncharacterized pupal eye and wing enhancer for the critical mitotic regulator Cdc25c (*string*) and establish that Mi-2 and E93 synergistically regulate the activity and shut-off of this enhancer *in vivo*. Our work on E93 and Mi-2 provides a long-missing molecular pathway linking ecdysone signaling in the pupa to the establishment and maintenance of the postmitotic state and reveals how this is coordinated with the progression of terminal differentiation.

## Results

### Mi-2 is required for stable cell cycle exit in the *Drosophila* eye and wing

Among *Drosophila* cell cycle genes, three key rate-limiting genes, *e2f1*, *cyclin E* and *string,* have uniquely complex modular cis regulatory elements that span ten to >50 kilobases, which control their tissue specific and temporal expression patterns [19–23]. E2F1 is a transcriptional activator that works with its binding partner DP to regulate the expression levels of hundreds of cell cycle genes across cell cycle phases and controls the overall timing of the cell cycle [24–27]. Cyclin E is an activating partner for the S-phase cyclin-dependent kinase and a rate-limiting factor that regulates the G1-S phase transition, and String (Cdc25c in mammals) is a phosphatase that removes an inhibitory phosphorylation on the mitotic Cyclin/Cdk complex and is rate-limiting for entry into mitosis in many *Drosophila* tissues [7, 26, 27]. We previously examined chromatin accessibility dynamics in the *Drosophila* wing, eye, and brain during metamorphosis, observing thousands of accessibility changes coordinated with morphogenesis, cell cycle exit and terminal differentiation [13, 14]. We found that enhancers at these three specific rate-limiting cell cycle genes undergo decommissioning after cell cycle exit, while the vast majority of other cell cycle genes exhibit maintained chromatin accessibility, even after prolonged exit from the cell cycle. Furthermore, in the wing, we showed that the elements at these three rate-limiting cell cycle genes continue to close, even in tissues subject to genetically driven, ectopic Cyclin/Cdk-induced proliferation at stages of metamorphosis when they would normally be postmitotic [13]. This demonstrates that the closing of these elements in the wing is developmentally controlled and independent of the tissue’s proliferative status. Based on these findings we developed a model where chromatin decommissioning at three rate-limiting cell cycle genes, which we hypothesize to be critical to maintain a robust non-cycling state, is mechanistically tied to the commencement of the terminal differentiation process via a temporal developmental cue.

To better understand the molecular mechanism(s) that mediate these chromatin accessibility changes at loci encoding key cell cycle regulators, we first sought to identify the chromatin remodeler(s) responsible. Given that enhancer regions at the loci of interest undergo similar postmitotic decommissioning in the wing, eye and brain [14], we reasoned that we could utilize adult eye size as a convenient phenotypic readout in an RNAi-based candidate screen to identify chromatin regulators that are required for cell cycle exit. We previously identified new molecular regulators of cell cycle exit using a sensitized genetic background where specific cell types in the *Drosophila* eye undergo 1-2 extra rounds of cell division. This is achieved with the overexpression of the G1-S regulator Cyclin E (CycE) driven by the eye-specific driver, *GMR*-Gal4 along with expression of P35, an apoptosis inhibitor [28]. This background generates enlarged adult eyes amenable to forward genetic or candidate screening for ‘enhancers’ that cause additional or prolonged proliferation and increase eye size, as well as ‘suppressors’ that reduce proliferation and decrease eye size [29]. As internal screening controls, we included a known suppressor condition, overexpression of the G1-S and G2-M Cyclin/Cdk inhibitors Dacapo and Wee, as well as a positive control for enhanced proliferation, overexpression of the positive cell cycle regulators Cyclin D, Cdk4 and E2F1. Among the screen’s positive hits, we focused on the chromatin remodeler Mi-2, because both Mi-2^RNAi^, as well as loss of one allele (*Mi-2^4^*/+), resulted in reproducibly larger eyes (Figure 1A). We stained pupal eyes from each of these conditions for cell boundaries to visualize whether the enlarged eyes resulted from increases in cell number. As expected, suppression with Dacapo + Wee overexpression reduced interommatidial cell density and cone cell numbers, while Mi-2^RNAi^ elevated interommatidial cell density and cone cell numbers, indicating that additional cell cycles had occurred with knockdown of Mi-2 (Figure 1B). Given the similarities in chromatin dynamics at cell cycle genes across tissues, we next examined whether Mi-2 also plays a role in cell cycle exit in pupal wings. Specifically, to test whether inhibition of Mi-2 further delays cell cycle exit in wings sensitized by ectopic proliferation, as it did in the eye screen, we co-overexpressed E2F1 and its partner DP (hereafter referred to as E2F complex) in the posterior pupal wing using *engrailed-*Gal4 with a temperature sensitive Gal80 repressor Gal80^TS^ (*en^TS^)*. We have previously observed that E2F expression in the wing increases nuclear density by driving additional cell cycles from 24-36h APF (hours after puparium formation) but cannot sustain cell cycling at later timepoints [9]. Here we find that co-expression of E2F+ Mi-2^RNAi^ allows for continued mitotic cycling at 42h APF (Figure 1C), demonstrating that in the wing Mi-2 also plays a role in promoting cell cycle exit at later timepoints under conditions where cell cycle exit has been delayed. Finally, we examined whether inhibition of Mi-2 in a non-sensitized background affects cell cycle exit in each of these tissues. We expressed a control RNAi (*white^RNAi^*) or *Mi-2*^RNAi^ in eyes during the final cell cycle using *GMR*-Gal4. Eyes expressing the control RNAi exhibit very few mitoses at 24h APF and little E2F activity as measured by expression of an E2F1 transcriptional reporter, PCNA-GFP [30], while expression of Mi-2^RNAi^ increases E2F activity and mitoses (Figure S1A). Overexpression of Mi-2^RNAi^ alone in the posterior pupal wing using *en^TS^*to activate expression beginning at L3 delays cell cycle exit and causes additional mitoses at 28h APF that resolve by 36h APF (Figure S1B). Thus, Mi-2 is a modulator of cell cycle exit timing in both the eye and the wing.

**Figure 1:**
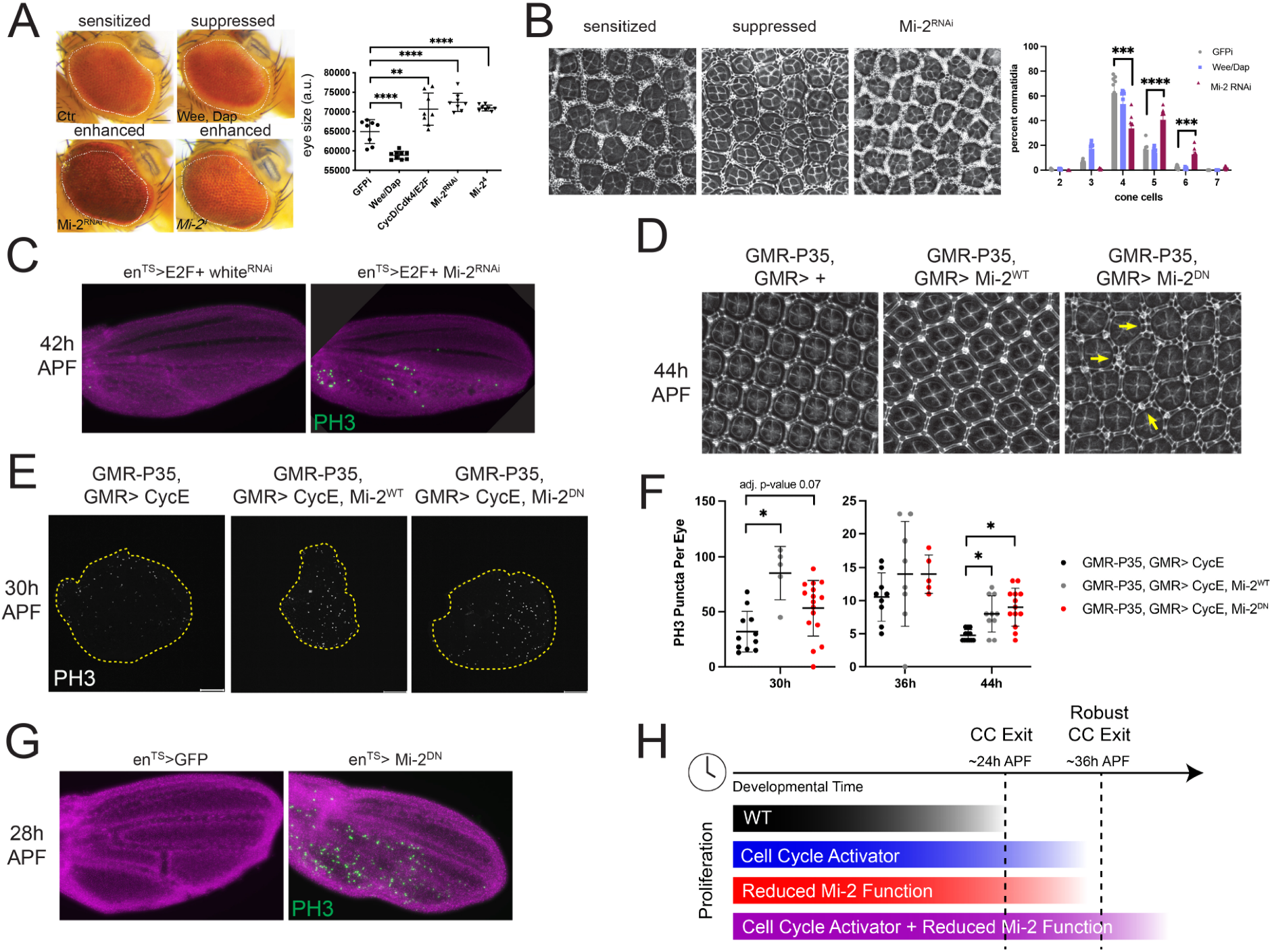
Mi-2 is required for stable cell cycle exit in the *Drosophila* eye and wing. (**A**) Representative images (left) and eye area quantifications (right) of adult eyes from a sensitized screen to identify chromatin remodelers that impact cell cycle exit. Dashed lines indicate the sensitized control eye size. Sensitized eyes express Cyclin E and a control RNAi (GFPi) under the control of *GMR*-Gal4, along with GMR-P35 to inhibit apoptosis. Controls include addition of cell cycle inhibitors Dacapo and Wee or positive cell cycle regulators CycD/Cdk4 + E2F. Experimental manipulations to inhibit Mi-2 function in the sensitized background include RNAi or a loss of function allele (*Mi-2^4^/+)*. Asterisks mark pair-wise comparisons with statistically significant differences via Mann-Whitney test. (**B**) Representative images (left) and cone cell number quantifications (right) from pupal eyes (36-40h APF) stained for cell boundaries with antibody against the septate junction protein Discs Large (Dlg). Sensitized, suppressed, and Mi-2^RNAi^ conditions as described for (A). Asterisks mark comparisons with statistically significant differences via pairwise t-tests. (**C**) 42h APF pupal wings overexpressing E2F (E2F1+DP) (left) or E2F + Mi-2^RNAi^ (right) in the posterior domain of the wing using temperature sensitive *engrailed*-Gal4 (*en^TS^*). DAPI to visualize DNA (magenta) and antibody recognizing phosphorylated Histone H3 (PH3) to visualize mitotic events (green). (**D**) 44h APF pupal eyes from control and eyes expressing wildtype Mi-2 (Mi-2^WT^) or dominant negative Mi-2 (Mi-2^DN^) stained for cell boundaries with antibody against Dlg. Yellow arrows indicate regions with ectopic interommatidial cells. (**E**) PH3 staining marking mitotic events in 30h APF pupal eyes from sensitized background and eyes additionally expressing Mi-2^WT^ or Mi-2^DN^. (**F**) Quantification of PH3 counts as in (E) at 30h, 36h and 44h APF. Asterisks mark comparisons with statistically significant differences; p-values derived from unpaired t-tests and corrected for multiple testing via Bonferroni-Dunn method. (**G**) 28h APF pupal wings overexpressing GFP (left) or Mi-2^DN^ (right) in the posterior domain of the wing using *en^TS^*. DAPI to visualize DNA (magenta) and antibody recognizing PH3 to visualize mitotic events (green). (**H**) Schematic diagram summarizing the role for Mi-2 in enforcing cell cycle exit in the pupa. Loss of Mi-2 activity alone results in delayed cell cycle exit relative to wildtype (WT) timing, and loss of Mi-2 can further extend proliferation in a sensitized background expressing a cell cycle activator such as CycE or E2F.

Mi-2 is a component of multiple chromatin remodeling complexes such as the NuRD and dMec complexes, where it acts enzymatically as an ATPase to power the movement of nucleosomes [31–33]. In addition, Mi-2 can form NuRD-independent complexes where the role of the dMec component MEP-1 is unclear [15, 34, 35]. To test whether the enzymatic ATPase activity of Mi-2 is required for proper cell cycle exit, we expressed a dominant-negative form of Mi-2 (K761R; Mi-2^DN^) in which the ATPase domain has been mutated; it has been shown that this protein cannot hydrolyze ATP [36]. Overexpression of Mi-2^DN^ but not wild type Mi-2 (Mi-2^WT^) in eyes resulted in ectopic interommatidial cells indicative of extra cell cycles in the pupal eye, demonstrating that the ATPase activity of Mi-2 is required for proper cell cycle exit (Figure 1D). Interestingly, in the CycE- and P35-expressing sensitized background we observed additional mitoses with the addition of either Mi-2^WT^ or Mi-2^DN^ at 30h and 44h APF (Figure 1E-F). However, we noted that despite its ability to induce more mitotic events at these pupal timepoints, Mi-2^WT^ ultimately suppressed adult eye size in the sensitized screening background (Figure S1C) suggesting overexpression of Mi-2^WT^ slows the cell cycle and reduces the overall amount of proliferation, perhaps due to altered stoichiometry of Mi-2 complexes involved in DNA damage repair and chromosome condensation [37, 38]. Finally, as in the eye, Mi-2^DN^ expressed during the final cell cycle under the control of *en^TS^*, is sufficient to delay cell cycle exit in the pupal wing (Figure 1G), demonstrating that the ATPase activity of Mi-2 is required for proper cell cycle exit in the *Drosophila* pupal eye and wing. Thus, Mi-2 plays a role in ensuring proper cell cycle exit during metamorphosis, both during normal cell cycle exit and in the context of genetically driven cell cycle exit delay (Figure 1H).

### Mi-2^DN^ expression compromises developmentally dynamic changes in chromatin accessibility

Given our finding that Mi-2 expression and enzymatic activity are required for proper timing of cell cycle exit in the pupal eye and wing, we next looked to identify the chromatin regions and gene transcripts dependent on Mi-2 for proper developmental dynamics. To this end, we performed ATAC-seq and RNA-Seq in pupal eyes, using the GMR-Gal4 driver to manipulate Mi-2 throughout the eye during the final cell cycle and as terminal differentiation processes are underway. We performed sample preparations at 24h APF, when control eyes are undergoing cell cycle exit, but Mi-2 manipulations cause ectopic cycling, and at 44h APF, when all of the tissues are postmitotic. We compared four conditions, each of which includes the apoptosis inhibitor P35: GMR-Gal4 driver alone (Control) and *GMR*-Gal4 driving Mi-2^RNAi^, Mi-2^WT^ or Mi-2^DN^. To begin to assess these datasets, we first performed multi-dimensional scaling analyses of all ATAC-seq and RNA-seq samples. Notably, we observed that the Mi-2^RNAi^ ATAC-Seq and RNA-Seq data clustered with the control condition at both 24h and 44h APF (Figure S2). Consistent with this, we saw variable and mild knockdown of the Mi-2 transcript in the Mi-2^RNAi^ RNA-Seq data, suggesting the Mi-2 knockdown was inconsistent. Because of this we did not analyze the Mi-2^RNAi^ samples further. Furthermore, we noted that by 44h APF the expression of Mi-2^DN^ or Mi-2^WT^ drive divergent transcriptomic changes as evidenced by these RNA-Seq samples clustering away from controls and away from each other, while only Mi-2^DN^ induced chromatin accessibility changes that are strongly divergent from control at the 44h APF time point (Figure S2). This suggests Mi-2^WT^ expression has effects on gene expression that are not reflected in coordinated chromatin accessibility changes. Because the samples overexpressing Mi-2^DN^ exhibit the most deviation from control samples in chromatin accessibility and gene expression at 44h APF, we chose to focus our further analyses primarily on these samples.

To validate our Mi-2^DN^ dataset, we first examined genes previously shown to be regulated by Mi-2 in other tissues and organisms. Previous work knocking down Mi-2 in *Drosophila* larval brains using a neuronal driver [32] reported ectopic expression of genes not typically expressed in the neuronal lineage; we observe that many of the top upregulated genes in this model are affected in a similar manner by Mi-2^DN^ expression in the eye (Figure S3A). Furthermore, we recapitulate the de-repression of germline genes, which was observed with Mi-2^RNAi^ in the brain [32] as well as in *C. elegans* where the homolog of Mi-2 has been shown to repress germline genes in somatic tissues [39] (Figure S3B). These data further support the interpretation that catalytically deficient Mi-2^DN^ interferes with the activity of the endogenous protein and phenocopies Mi-2 loss of function.

Comparing chromatin accessibility data from eyes expressing Mi-2^DN^ to controls, we identified 2,284 differentially accessible regions at the 44h APF time point, with the majority of these regions exhibiting increased accessibility when Mi-2 function is compromised (n=1,893). (Figure 2A). Motif enrichment analysis of the differentially accessible regions relative to unaffected regions identified binding motifs for many classes of transcription factors, including homeodomain, bZIP and C2H2 zinc finger families; notably, many of these are factors previously implicated as regulators of eye development, including Glass, Rough, and Pph13 (Figure S3C and Table S1) [40–43]. When we further assessed differential motif enrichment by testing regions that gained accessibility with Mi-2^DN^ against those that showed reduced accessibility, we found that a number of these motifs are differentially enriched between these groups (Figure S3C and Tables S2-3), suggesting that Mi-2’s remodeling action at chromatin- whether to reduce or enhance accessibility- may be influenced by the transcription factors present at its target regions. Overall, these results support the idea that as a general chromatin remodeler, Mi-2 likely cooperates with a diverse variety of developmental factors to modulate chromatin accessibility in the developing eye.

**Figure 2:**
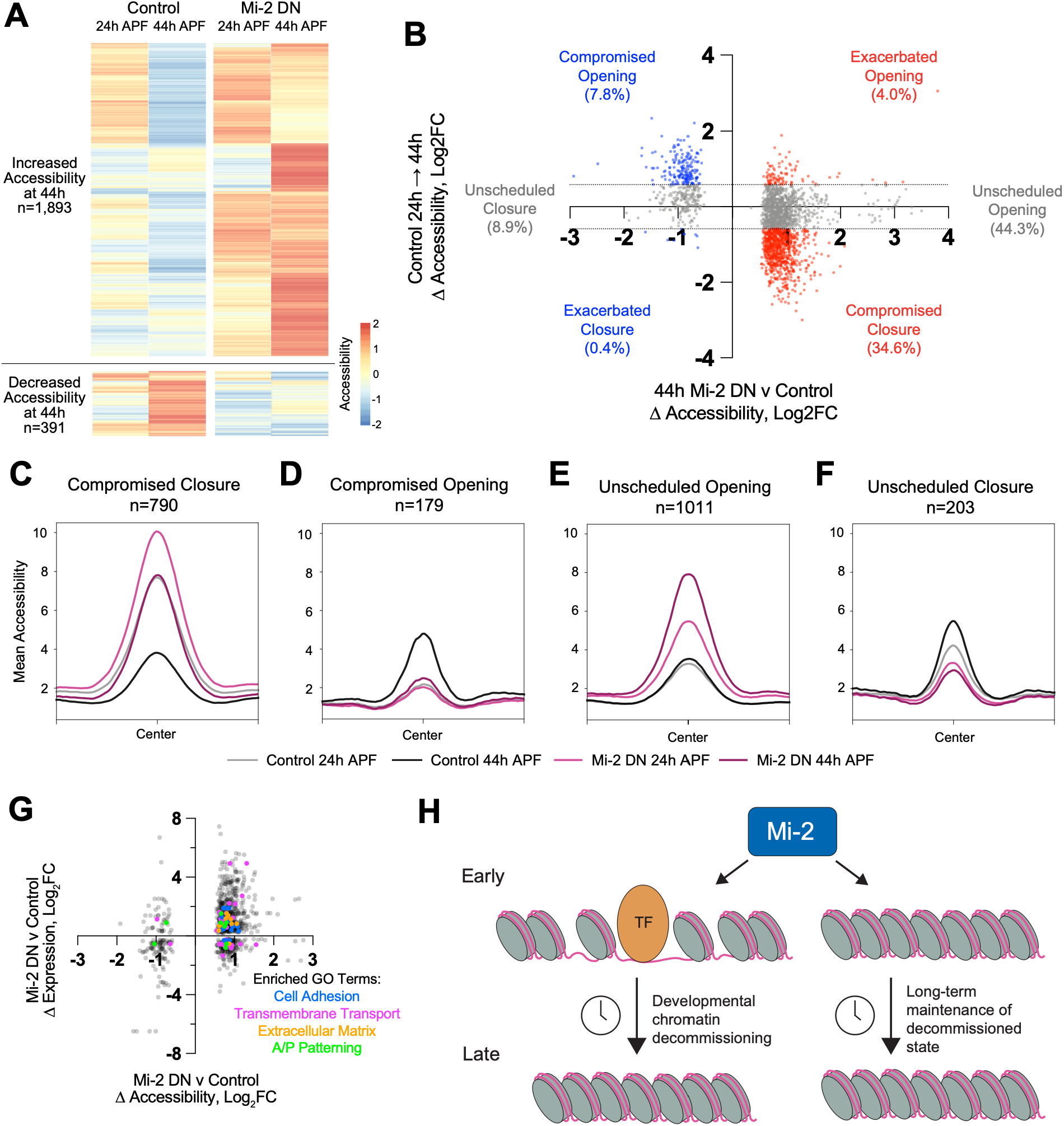
Mi-2 limits chromatin accessibility at developmentally dynamic and inaccessible regions in the developing eye. (**A**) Heatmap showing ATAC-Seq chromatin accessibility measures from control and Mi-2^DN^-expressing eyes at regions that exhibit differential accessibility with Mi-2^DN^ expression relative to control 44h APF. ATAC-Seq coverage values are scaled by Z-score and regions are hierarchically clustered. (**B**) Scatterplot showing the relationship between the change in accessibility induced by Mi-2 DN expression and the change in accessibility for the same regions in the control eye time course from 24h to 44h APF. Regions with Log2FC values between −0.585 and 0.585 in the control time course were considered stable and are colored in grey. All other points are colored red (increased accessibility with Mi-2 DN) or blue (decreased accessibility with Mi-2 DN). (**C**-**F**) Line plots showing the average accessibility measures (count-per-million normalized read coverage) for regions falling into the most prominent dynamics profiles as indicated in panel B. Plots include samples at 24h (control: grey, Mi-2 DN: light pink) and 44h (control: black, Mi-2 DN: dark pink); accessibility was averaged in 10 base pair bins over a 1 kilobase region centered on the peak center. (**G**) Scatterplot showing the relationship between Mi-2^DN^-induced chromatin accessibility (x-axis) and gene expression changes (y-axis) at 44h APF, for all genes with differential expression that are also associated with differentially accessible chromatin regions. Colored dots highlight data points from genes that fall within significantly enriched GO Terms as listed next to the plot. (**H**) Model describing the predominant roles of Mi-2 in terminally differentiating tissues.

By comparing the changes in accessibility induced by Mi-2^DN^ to the accessibility dynamics that the same regions show in control samples from 24h to 44h APF, we note a few prominent profiles among the regions responsive to Mi-2^DN^ (Figure 2B and Figure S4). First, compromising Mi-2 function opposes the developmental dynamics that occur at many regions over the 24h to 44h time course, either by compromising closure/decommissioning (34.6% of regions; Figure 2B lower right and Figure 2C) or compromising opening (7.8% of regions; Figure 2B upper left and Figure 2D). These findings indicate an apparent slowing or failure of developmental remodeling with Mi-2^DN^, suggesting that Mi-2 directly or indirectly orchestrates accessibility changes at developmentally dynamic regions of the genome. This is consistent with its known enzymatic activity and with the idea that Mi-2 is important for cell state transitions. Further, we note another subset of regions that exhibit little to no accessibility change in the control time course but where Mi-2^DN^ alters accessibility, predominantly by inducing what we refer to as an “unscheduled opening” (44.3% of regions; Figure 2B middle right and Figure 2E) or to a lesser extent “unscheduled closure” (8.9% of regions; Figure 2B middle left and Figure 2F). We wondered whether the large group of regions undergoing an unscheduled opening in the presence of Mi-2^DN^ might be comprised of regulatory regions that were utilized earlier in eye development. However, in our previously generated control time course data we see that these regions exhibit very little to no accessibility even at the earlier L3 time point (Figure S5A); given the limited resolution of this time course, it remains possible that this group includes regulatory regions that are accessible in the eye at some other developmental time point and are temporally misregulated in the presence of Mi-2^DN^. However, given that Mi-2 has previously been implicated in the repression of ectopic gene expression in other tissues, this group could also represent accessibility gains at regulatory regions typically only utilized in other lineages. As a whole, these data imply that in addition to its role in mediating developmental dynamics, Mi-2 plays an essential role in the maintenance of inaccessible chromatin in the pupal eye.

To understand how Mi-2 inhibition impacts gene expression in the developing eye, we intersected these chromatin accessibility data with gene expression data by assigning chromatin regions to the nearest gene and found that 39% of chromatin regions with altered accessibility upon Mi2^DN^ expression reside near a gene that is differentially expressed. Most of the differential accessibility-differential gene expression pairings reflect gain of accessibility associated with increased gene expression in Mi2^DN^ eyes compared to control (Figure 2G), and many of these target genes show developmentally dynamic expression in the control time course that is compromised by Mi-2^DN^ (Figure S5). Assessing the types of genes that reside near differentially accessible regions, we find enrichments of genes associated with developmentally relevant terms including tissue patterning, cell adhesion and extracellular matrix functions, all of which are known to be important processes in the early pupal stages of eye development (Figure 2G, Table S4). These findings suggest that Mi-2^DN^ expression may compromise the differentiation of the eye; this is further supported by gene ontology enrichment analysis of all downregulated genes, where differentiation-associated terms are enriched, including visual perception and intracellular signaling (Table S5). As a whole, our genome-wide surveys of accessibility and transcriptional changes caused by Mi-2^DN^ in the eye are consistent with the idea that Mi-2 is directly or indirectly responsible for modulating accessibility at dynamic regulatory regions to support developmental gene expression programs. In particular, Mi-2 appears to contribute to the developmental progression of this tissue predominantly by mediating decommissioning of dynamic regulatory regions and through maintenance of the inaccessible chromatin state to prevent inappropriate gene expression during terminal differentiation (Figure 2H).

### Mi-2^DN^ alters transcript expression and chromatin accessibility at cell cycle genes

We initially identified Mi-2 as a strong candidate to modulate chromatin dynamics contributing to cell cycle exit. Therefore, a pressing question was to determine the effect of Mi-2^DN^ expression at cell cycle gene loci and particularly the chromatin dynamics at complex, rate-limiting cell cycle genes. In addition to our earlier observations that Mi-2^DN^ expression compromises cell cycle exit, our gene ontology analyses of upregulated genes at 44h APF identified a significant enrichment of genes associated with DNA repair (Table S5), suggesting persistent expression of factors involved in DNA synthesis and quality control. We next assessed the overall expression of ∼300 curated cell cycle genes from control and Mi-2^DN^ samples. In controls most of the cell cycle genes undergo transcriptional repression as expected from 24h APF, when the tissue exits the cell cycle, to 44h APF when the tissue has been postmitotic for 20 hours and differentiation is well underway (Figure 3A). In Mi-2^DN^ expressing eyes, many of the cell cycle gene transcripts are expressed at higher levels than control at both the 24h and 44h APF time points. These data support the idea that Mi-2 is important for the transition from proliferative activation to postmitotic transcriptional repression of the cell cycle program. We next asked whether expression of the three rate-limiting cell cycle genes that show complex postmitotic chromatin dynamics, *cycE*, *e2f1* and *string (stg)*, have altered expression with Mi-2^DN^. *cycE* expression is not altered by Mi-2^DN^ (Figure 3B), but Mi-2^DN^ expression slightly changes the expression levels of *e2f1* (Figure 3C, 114% of control at 24h APF and 81% of control at 44hAPF) and modestly elevates *string* expression (Figure 3D, 138% and 132% of control levels at 24h and 44h APF, respectively). Interestingly, both *e2f1* and *string* show chromatin accessibility changes with Mi-2^DN^ expression (Figure S6). We were particularly intrigued by the change in accessibility we observed with Mi-2^DN^ at a region far upstream of *string*; this region shows prominent dynamics in our previous time course studies with a peak of accessibility at 24h APF followed by near-complete decommissioning at the 44h time point (Figure 3E). With Mi-2^DN^, the decommissioning of this element is compromised at 44h APF (Figure 3F). Notably, this region is in close proximity but distinct from a known larval eye enhancer region for *stg*, called *stg*-Visual System (*stg*-VS) [23]. Together with the change in *string* transcript levels, the persistent accessibility at this element suggests that Mi-2 may exert its effects on the proliferative status of the tissue in part by modulating the postmitotic chromatin dynamics at an uncharacterized candidate regulatory element for *string*, a critical, rate-limiting mitotic factor.

**Figure 3:**
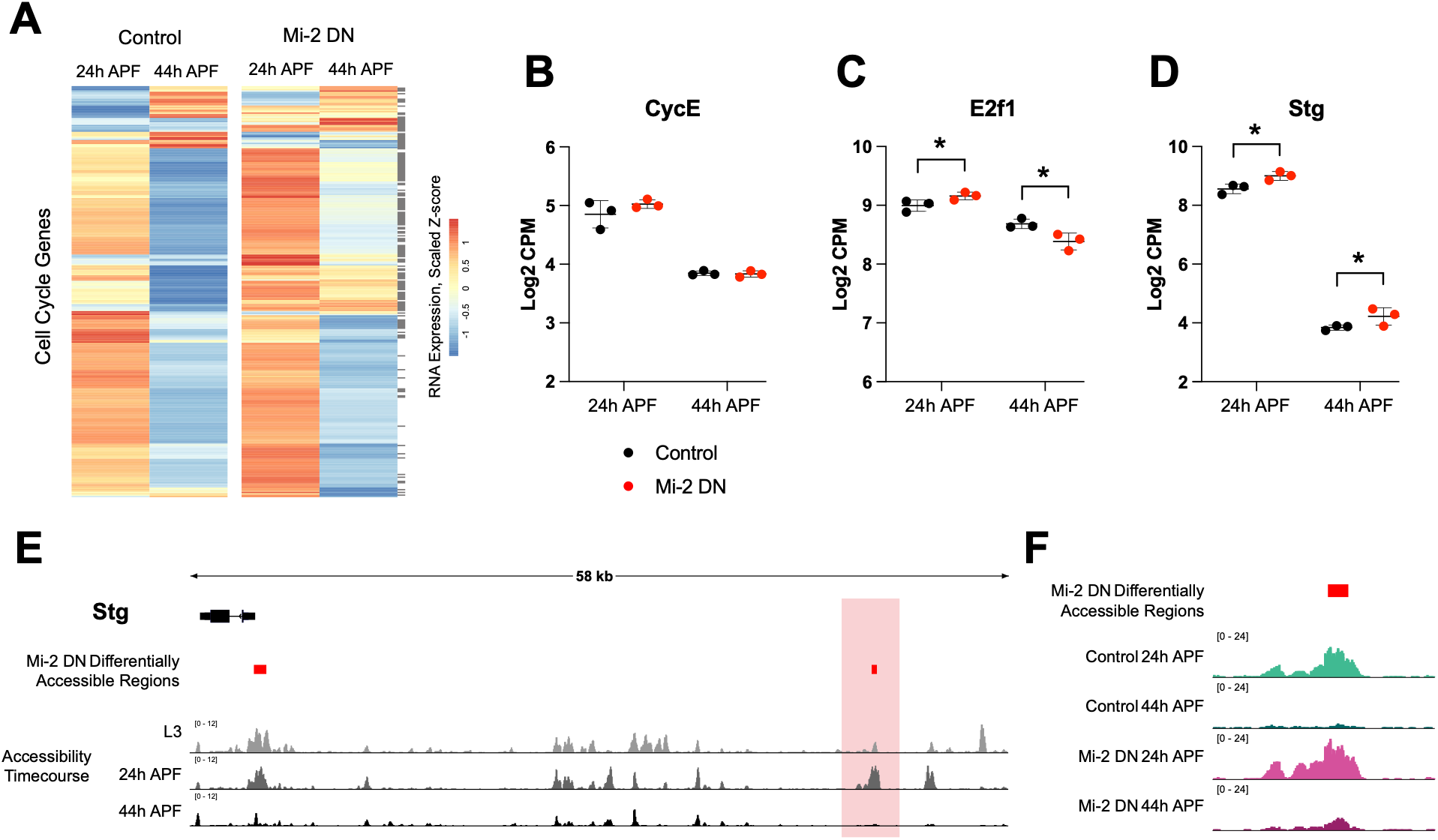
Mi-2^DN^ compromises postmitotic repression of cell cycle genes and chromatin decommissioning at the mitotic regulator String. (**A**) Heatmap showing average gene expression values for cell cycle genes in control and Mi-2^DN^-expressing eyes at 24h and 44h APF. Gray boxes at the right side of the heatmap mark genes that are differentially expressed in Mi-2^DN^ versus control at 24h or 44h APF. Gene expression values are plotted as scaled Z-scores and are hierarchically clustered. (**B-D**) Gene expression values from independent RNA-Seq replicates for *Cyclin E* (B, CycE), *E2f1* (C), and *string* (D, Stg) in control (black) and Mi-2^DN^ expressing (red) eyes at 24h and 44h APF. Asterisks mark statistically significant comparisons with FDR-corrected p-value less than 0.05. (**E**) Diagram of the gene locus encoding Stg with RefSeq transcript indicated in black. Data tracks depict ATAC-Seq data from a time course study of eyes at L3, 24h and 44h APF. Y-axes indicate count-per-million normalized read coverage. Regions that exhibit differential accessibility in Mi-2^DN^ relative to control at 44h APF are indicated with red boxes. (**F**) View of the region highlighted with the light red box in panel E, showing ATAC-Seq data from control and Mi-2^DN^ expressing eyes at 24h and 44h APF. Y-axes indicate count-per-million normalized read coverage.

### Mi-2^DN^ compromises the ecdysone temporal cascade

Our observation that Mi-2^DN^ expression compromises temporal chromatin dynamics (Figure 2) and differentiation-associated gene expression programs (Tables S4-S5) raised the question of whether Mi-2 may support timely cellular responses to developmental cues during metamorphosis. The best characterized such developmental cue in *Drosophila* is ecdysone, a systemic hormone that is known to trigger organismal transitions throughout development including terminal differentiation events during metamorphosis. Importantly, it has long been established that a large systemic pulse of high ecdysone titers occurs beginning around 24h APF, and this pulse has been implicated in the differentiation of adult tissues [44, 45]. To assess whether Mi-2 may play a role in the ecdysone response in the developing eye during this time frame, we assessed the transcript expression levels for several previously characterized ecdysone response genes in control and Mi-2^DN^ samples. Many of these transcripts are expressed in a time point-specific manner in the controls, wherein ‘early’ targets expressed highly at 24h APF are largely repressed by 44h APF while distinct ‘late’ target genes are specifically expressed at 44h APF (Figure 4A). These patterns are consistent with the Ashburner model for the ecdysone temporal cascade, wherein after a cell receives the hormonal signal early target gene expression is induced, followed by the early products inducing late targets while also repressing their own expression to generate a sequential cascade [46]. Interestingly, Mi-2^DN^ expression causes an apparent slowing or stalling of the cascade, as indicated by persistent 44h APF expression of early transcripts and reduced induction of late targets. Notable early target genes showing persistent expression with Mi-2^DN^ are *Broad* (Figure 4B, 180% of control at 44h APF), *Blimp-1* (Figure 4C, 210% of control at 44h APF) and *Eip93F* (Figure 3D, 170% of control at 44h APF), and an example of a late target with compromised expression is *Eip78C* (Figure 4E, 35% of control at 44h APF). Each of these genes exhibit chromatin accessibility changes with Mi-2^DN^ alongside their expression changes (Figure S7). Of particular interest are the expression and accessibility changes we observe at *Broad* and *Blimp-1* (Figure 4F-I). Each of these genes has been previously described to play roles in the developing eye and their misregulation further supports the idea that differentiation pathways are compromised with Mi-2^DN^ expression [47, 48]. Taken together, these findings support the idea that Mi-2 directly or indirectly contributes to the chromatin dynamics mediating the ecdysone response during metamorphosis, which is likely to at least partially explain the differentiation defects observed with Mi-2^DN^ expression.

**Figure 4:**
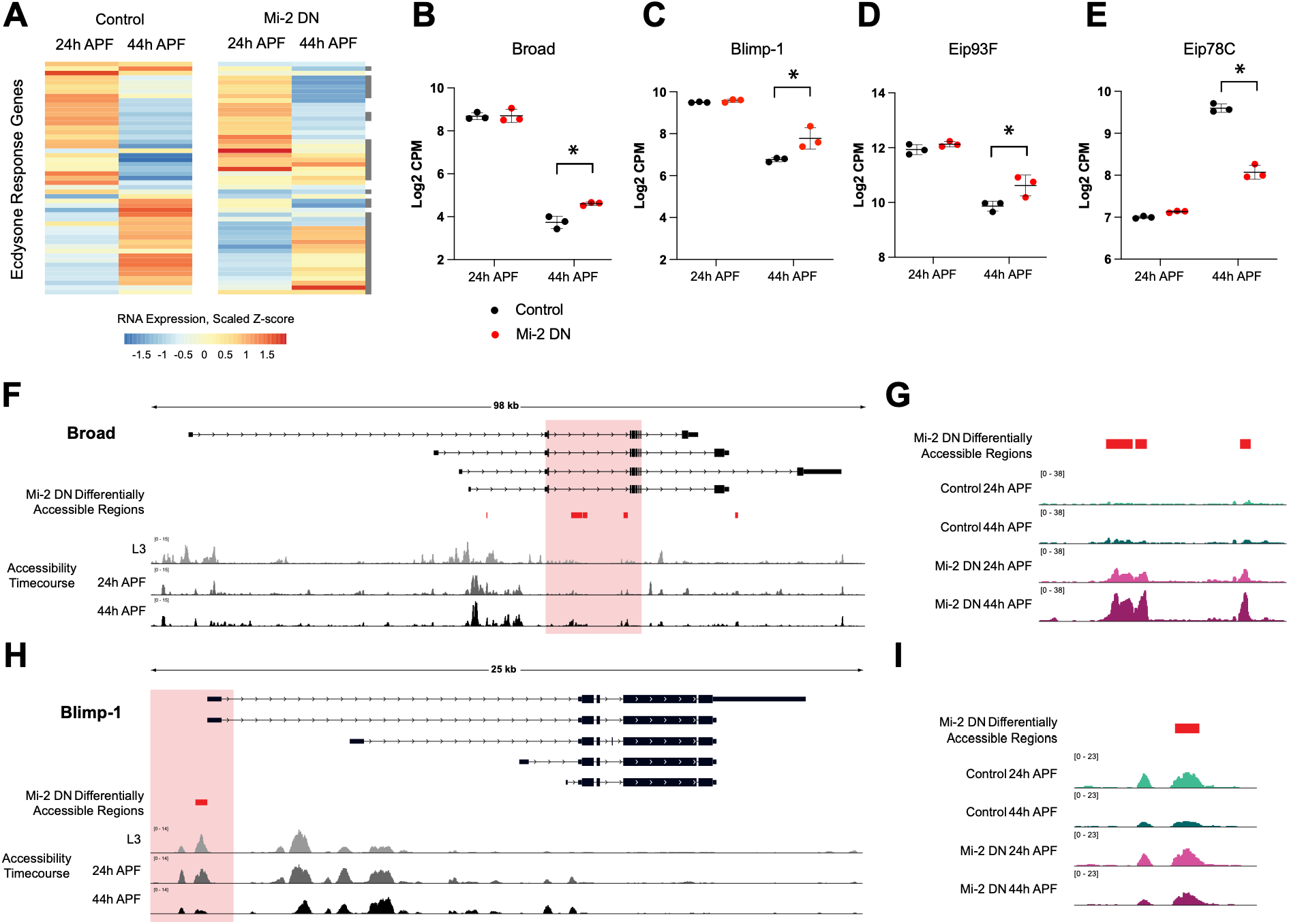
Mi-2^DN^ compromises the ecdysone temporal cascade. (**A**) Heatmap showing average gene expression values for ecdysone response genes in control and Mi-2^DN^ expressing eyes at 24h and 44h APF. Gray boxes at the right side of the heatmap mark genes that are differentially expressed in Mi-2^DN^ versus control at 24h or 44h APF. Gene expression values are plotted as scaled Z-scores. (**B-E**) Gene expression values from independent RNA-Seq replicates for *broad* (B), *Blimp-1* (C), *Eip93F* (D), and *Eip78C* (E) in control (black) and Mi-2^DN^ expressing (red) eyes at 24h and 44h APF. Asterisks mark statistically significant comparisons with FDR-corrected p-value less than 0.05. (**F**,**H**) Diagrams of the gene loci encoding *broad* (F) and *Blimp-1* (H) including simplified summaries of RefSeq-annotated transcripts, with arrows indicating the direction of transcription. Data tracks depict ATAC-Seq data from time course series of eyes at L3, 24h and 44h APF. Y-axes indicate count-per-million normalized read coverage. Regions that exhibit differential accessibility in Mi-2^DN^ relative to control are highlighted in red. (**G**,**I**) Views of the regions at *Broad* (G) and *Blimp-1* (I) highlighted with the light red boxes in panels F and H, showing ATAC-Seq data from control and Mi-2^DN^ expressing eyes at 24h and 44h APF. Y-axes indicate count-per-million normalized read coverage.

### Mi-2 and E93 regulate accessibility at a subset of overlapping dynamically accessible regions

General chromatin remodelers such as Mi-2 are thought to be recruited to specific genomic locations at the right times during development through interactions with sequence-specific transcription factors [34, 35]. Indeed, Mi-2 is expressed in the eye throughout the time course from L3 to 44h APF, suggesting that there must be some factors it interacts with to mediate its temporally specific effects on chromatin accessibility and gene expression. Moreover, our previous finding that postmitotic decommissioning at complex cell cycle genes occurs on schedule despite genetically-driven ectopic proliferation supports the idea that there is a signal directing the timing of remodeling at these genes. The observation that Mi-2^DN^ expressing eyes exhibit compromised decommissioning at the *string* locus along with an apparent defect in the ecdysone transcriptional response suggested to us that there may be some direct or indirect interactions between Mi-2 and the cellular ecdysone response that could coordinate *string* enhancer decommissioning at the proper time during metamorphosis.

Despite its nature as a secreted hormone that circulates systemically, diverse cell types and tissues can vary widely in their response to the same ecdysone pulse [49, 50]. Downstream of its nuclear receptor (Ecdysone Receptor, EcR), the transcriptional response to ecdysone is mediated through a series of sequentially expressed transcription factors; the same set of ecdysone response factors are generally utilized in many tissues throughout the body, but tissue-specific ecdysone responses have been associated with modifications to the timing and sequence of expression of these factors [50]. If Mi-2 coordinates with a factor downstream of ecdysone to trigger cell cycle gene decommissioning in the eye and the wing, we would expect this factor to be expressed in both of these tissues at the 24h APF time point when cell cycle exit and subsequent cell cycle gene remodeling are set in motion. Using our previously generated RNA-Seq time course data from each of these tissues [6, 14], we compared expression of ecdysone-induced transcription factors in the wing and the eye at the L3, 24h and 44h APF time points. This revealed ecdysone response genes expressed with similar dynamics in the two tissues, such as the early gene *Broad* (*Br*) and late gene *Ecdysone induced protein 75B* (*Eip75B*) while others such as *Ecdysone induced protein 63E* (*Eip63E*) show tissue-specific expression dynamics (Figure 5A). However, one transcription factor exhibits a peak of transcript expression in both tissues at the 24h APF time point: *Ecdysone induced protein 93F* (*Eip93F; E93*). Indeed, using an endogenously tagged allele of *E93*, we confirmed that this protein is expressed at pupal stages in the eye and the wing (Figure S8A).

**Figure 5:**
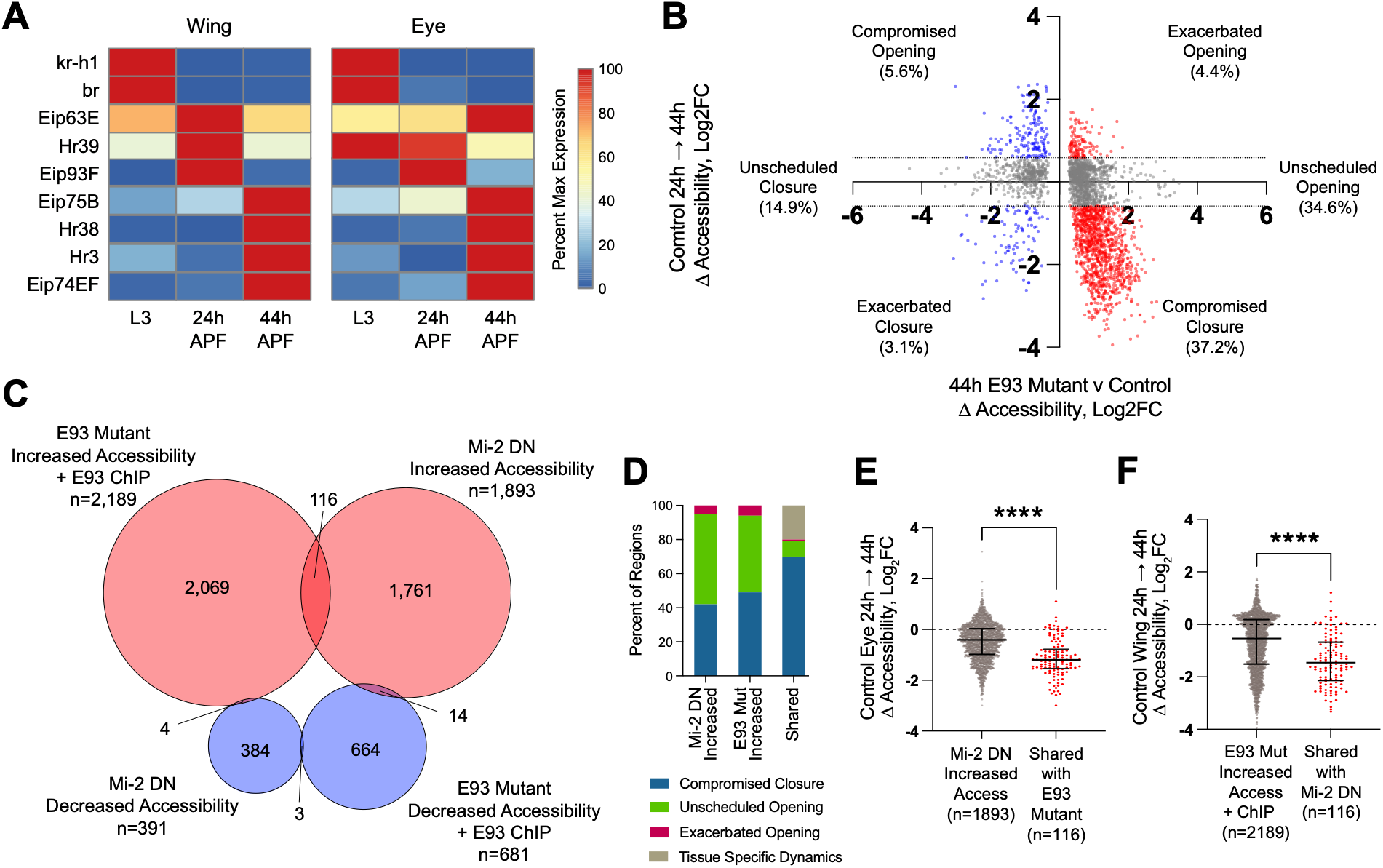
Mi-2 and E93 regulate accessibility at a subset of overlapping dynamically accessible regions. (**A**) Heatmap showing average gene expression values for a selection of ecdysone-induced transcription factors in a time course series of eyes at L3, 24h and 44h APF. Gene expression values are plotted as percent of maximum for each respective tissue. (**B**) Scatterplot showing the relationship between the change in FAIRE-Seq accessibility induced by loss of E93 in the 44h APF wing and the change in accessibility for the same regions in the control wing time course from 24h to 44h APF. Regions with Log2FC values between −0.585 and 0.585 in the control time course were considered stable and are colored in grey. All other points are colored red (increased accessibility in E93 mutant) or blue (decreased accessibility in E93 mutant). (**C**) Venn diagram showing the intersections of regions that are differentially accessible with Mi-2^DN^ expression in the 44h APF eye and regions that are differentially accessible with loss of E93 in the 44h APF wing and are bound by E93 in 24h APF wings. (**D**) Stacked bar plots showing the proportion of dynamic profiles among regions that have increased accessibility with Mi-2 DN in eyes (n=1893), have increased accessibility in E93 mutant wings and are bound by E93 (n=2189), and regions shared between the two datasets (n=116). Blue indicates the proportion that exhibit compromised closure, green indicates unscheduled opening, magenta indicates exacerbated opening, and grey indicates ‘shared’ regions that exhibit tissue specific dynamics. (**E**) Plot showing the change in accessibility in control eyes from 24h to 44h APF, among all regions with increased accessibility in Mi-2 DN eyes at 44h or among regions ‘shared’ with E93 mutant wings. Error bars indicate standard deviation and asterisks mark statistical significance, p<0.0001 in Welch’s t-test. (**F**) Plot showing the change in accessibility in control wings from 24h to 44h APF, among all regions with increased accessibility in E93 mutant wings at 44h or among regions ‘shared’ with Mi-2 DN expressing eyes. Error bars indicate standard deviation and asterisks mark statistical significance, p<0.0001 in Welch’s t-test.

E93 has previously been characterized as a driver of terminal differentiation events during the pupal stages [51, 52]. We previously showed that E93 is essential for chromatin accessibility changes during wing metamorphosis, including opening and closing of chromatin [6]. However, it has remained unclear which chromatin remodelers are recruited by E93 to modify chromatin accessibility. Indeed, we recently determined that the *Drosophila* SWI/SNF chromatin remodeling complex is dispensable for mediating changes in chromatin accessibility in pupal wings despite contributing to E93-dependent gene regulation [53]. To identify potential interacting partners of E93 we performed a proximity labeling assay followed by mass spectrometry. Briefly, we generated stable *Drosophila* S2 cell lines in which E93 fused to a hyperactive biotin ligase (TurboID) is under inducible control. Mass spectrometry analysis of affinity purified proteins identified Mi-2 among the top hits (Table S6 and S7). Confidence in the quality of the proximity ligation results was supported by the presence of the transcriptional co-repressor, CtBP, among the top hits; prior studies have demonstrated interactions between CtBP and the E93 ortholog, LCoR, in breast cancer cells through its PxDLS recognition motifs, which are conserved in E93 [54]. Notably, CtBP is a member of the TORC chromatin remodeling complex along with Tou, which was also identified as a top hit in our E93 TurboID experiments. Altogether, these findings suggest that E93 and Mi-2 may interact, providing a possible mechanism through which Mi-2’s remodeling activity may be targeted to particular genomic locations during this critical developmental stage where cell cycle exit and terminal differentiation are coordinated.

Given that we have previously measured chromatin accessibility in the E93 mutant pupal wing and that we consider the processes governing cell cycle decommissioning likely to be shared between wing and eye, we hypothesized that comparing chromatin accessibility profiles across these two tissues would reveal candidate sites of coordinated regulation by Mi-2 and E93, including those that may be relevant to cell cycle exit. Furthermore, as we are primarily interested in identifying regions where E93 binds and may recruit and/or interact with Mi-2 to remodel chromatin, we incorporated E93 ChIP-Seq profiles from 24h APF pupal wings [6] to restrict the candidate target regions to sites of E93 binding. We reanalyzed our previously published accessibility data from control and *E93* mutant (*E93*^MUT^) pupal wings using the same scheme we applied to the accessibility data from Mi-2^DN^ expressing eyes (Figure 2C) and observed that many of the E93-bound regions with altered accessibility in *E93^MUT^*wings at 44h APF are developmentally dynamic in the control wing time course from 24h to 44h (Figure 5B). As with Mi-2^DN^ in the eye, the majority of regions responding to loss of E93 in the 44h wing gain accessibility, with most exhibiting a compromised closure (37.2% of regions; Figure 5B lower right) or an unscheduled opening (34.6% of regions; Figure 5B middle right).

To identify chromatin regions that may be coordinately regulated by Mi-2 and E93 we next intersected our 44h APF Mi-2^DN^ and *E93^MUT^* datasets and identified a limited number of candidate co-regulated or ‘shared’ regions (Figure 5C). The predominant intersection among these datasets are 116 regions that gain accessibility at 44h APF with Mi-2^DN^ and loss of E93, representing a significantly greater degree of overlap than predicted by chance (compared to 1000 permutations of randomized regions, Z=17.89, p<0.001) and suggesting that Mi-2 and E93 are most likely to work together to limit chromatin accessibility. Furthermore, assessing the dynamics of these regions revealed that they are significantly enriched for those that exhibit “compromised closure” in both Mi-2^DN^ expressing eyes and E93^MUT^ wings (Figure 5D, Two Proportion Z-tests: relative to Mi-2^DN^ increased accessibility regions Z=5.93 and p<0.001; relative to E93^MUT^ increased accessibility regions Z=4.41 and p<0.001). Consistent with this, the accessibility at these 116 ‘shared’ regions is significantly more developmentally dynamic from 24 to 44h APF compared to all regions that gained accessibility with Mi-2^DN^ in the eye (Figure 5E) or loss of E93 in the wing (Figure 5F). Finally, we note that by intersecting accessibility data with gene expression data from Mi-2^DN^ expressing eyes, the 116 shared regions reside near differentially expressed genes at a significantly higher rate compared to all regions with increased accessibility (54% versus 39%, Two Proportion Z-test: Z=3.27, p=0.001), suggesting that these regions are enriched for those that exert detectable effects on nearby gene expression. As a whole, these analyses identified a curated selection of candidate regions that Mi-2 and E93 may coordinately regulate across tissues and suggest that at shared target regions these factors predominantly act to decommission developmentally dynamic regulatory elements.

### E93 knockdown delays cell cycle exit and disrupts terminal differentiation in the pupal eye

While we have previously shown that E93 plays an important role in wing maturation, it is unknown whether E93 plays a role in eye development, and it has not previously been implicated in cell cycle exit in any context. To examine E93 function in eyes, we performed an RNAi knockdown of E93 using the *GMR*-Gal4 driver and P35 expression. The efficacy of E93 knockdown was confirmed by staining for loss of protein expression using the endogenously tagged E93 allele (Figure S8B). E93 knockdown with *GMR*-Gal4 results in obvious morphological and pigmentation defects at the adult stage, demonstrating that E93 indeed plays a role in eye development (Figure 6A). Although the adult E93^RNAi^ eyes appear smaller than controls, we note that the pupal eye disc is much thicker in the E93 knockdown condition (Figure S8C), suggesting that the thicker retina could arise from defects in cell cycle exit. We stained pupal eyes to assess cone cell and interommatidial cell boundaries but observed that the E93 knockdown pupal eyes have a highly disorganized structure with cell junctions that were difficult to resolve, such that we were unable to accurately assess cell number. However, staining for mitoses with E93^RNAi^, either in the CycE-expressing sensitized background (Figure 6B-C) or alone (Figure S8D), revealed a clear increase in mitotic events. Moreover, E93 knockdown in the wing also delayed cell cycle exit (Figure S8B), demonstrating that E93, like Mi-2, plays an essential role in eyes and wings to promote timely cell cycle exit.

**Figure 6:**
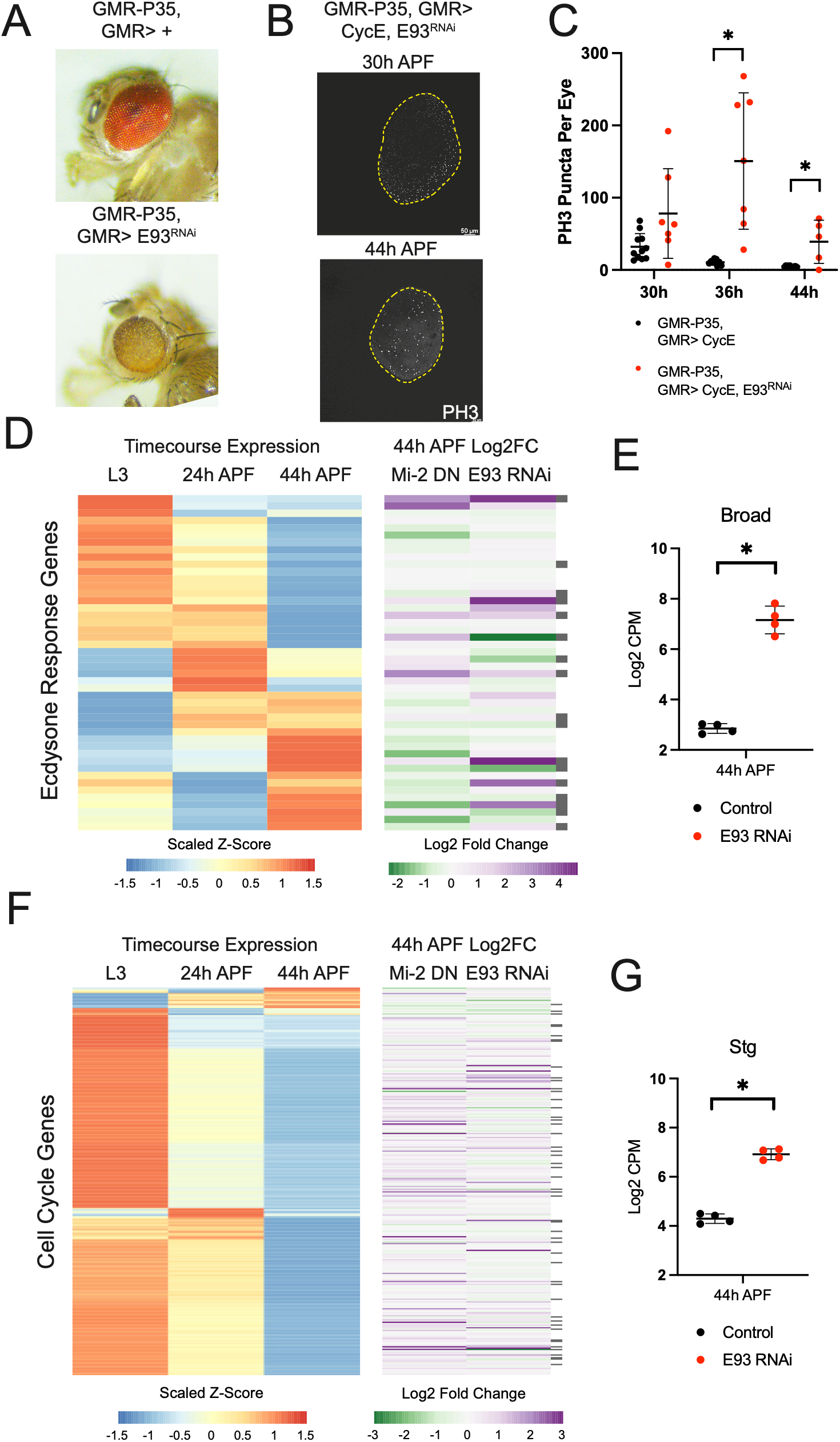
E93 knockdown delays cell cycle exit and disrupts terminal differentiation. (**A**) Representative images of adult eyes from GMR-P35, GMR-Gal4 control and with E93^RNAi^. (**B**) 30h and 44h APF eyes in the Cyclin E (CycE)-expressing sensitized background with expression of E93^RNAi^, stained with an antibody recognizing phosphorylated Histone H3 (PH3) marking mitotic events. (**C**) Quantification of PH3 puncta from 30h, 36h and 44h APF eyes as in (B). Note these PH3 experiments were performed in parallel with the Mi-2 data in Fig1F and therefore use the same CycE-expressing control for comparison. (**D**) Heatmap showing average ecdysone response gene expression levels from an RNA-Seq time course in control eyes at L3, 24h and 44h APF (left). Expression values are scaled by Z-score. To the right, a heatmap for the same genes, color coded by the log-transformed expression changes in 44h APF Mi-2^DN^-expressing eyes relative to their control, and in 44h APF E93^RNAi^ eyes relative to their control. Gray boxes to the right of the fold change heatmap mark the genes that showed differential expression in both datasets. (**E**) Gene expression values from independent RNA-Seq replicates for *broad* in control (black) and E93 RNAi-expressing (red) eyes at 44h APF. Asterisks mark statistically significant comparison with FDR-corrected p-value less than 0.05. (**F**) Heatmap showing average cell cycle gene expression levels from an RNA-Seq time course in control eyes at L3, 24h and 44h APF (left). Expression values are scaled by Z-score. To the right, a heatmap for the same genes, color coded by the log-transformed expression changes in 44h APF Mi-2^DN^-expressing eyes relative to their control, and in 44h APF E93^RNAi^ eyes relative to their control. Gray boxes to the right of the fold change heatmap mark the genes that showed differential expression in both datasets. (**G**) Gene expression values from independent RNA-Seq replicates for *string* in control (black) and E93 RNAi-expressing (red) eyes at 44h APF. Asterisks mark statistically significant comparison with FDR-corrected p-value less than 0.05.

To assess the transcriptional impact of E93 knockdown, we performed RNA-seq on eyes expressing E93^RNAi^ at 44h APF. As a member of the ecdysone transcriptional cascade, we assessed the expression of previously characterized ecdysone response genes in these eyes and found that E93^RNAi^ perturbs the transcriptional cascade, though in an apparently different manner than that of Mi-2^DN^. Rather than a clear bias toward persistent expression of 24h APF targets and muted induction of late targets, E93^RNAi^ caused more complex effects with early and late genes being variably down- and up-regulated (Figure 6D). This is consistent with the idea that E93 plays an instructive role to activate or repress its target genes in a locus-specific manner rather than acting as a general mediator of developmental transitions in the manner of Mi-2. As with Mi-2^DN^, E93 knockdown caused an increase in *Broad* transcript levels at 44h APF (Figure 6E). Given its ability to delay cell cycle exit, we also assessed cell cycle gene expression with E93^RNAi^ and observed persistent expression of many cell cycle genes at 44h APF (Figure 6F), consistent with a role in the postmitotic transcriptional repression of the cell cycle. Intriguingly, we noted increased expression of the mitotic regulator *string* with E93 knockdown as we had observed with Mi-2^DN^ (Figure 6G). Taken as a whole, phenotypic, chromatin accessibility and gene expression data from the eye and the wing suggest that E93 likely acts as a temporal factor to coordinate cell cycle exit with terminal differentiation. Based on the possible interaction between Mi-2 and E93 via proximity ligation and the available data identifying their genomic targets, we hypothesize that these two factors may operate in a coordinated manner at a small number of coregulated loci to coordinate the chromatin accessibility changes required for differentiation and cell cycle exit.

### Mi-2 and E93 regulate accessibility at a novel pupal *stg* enhancer after cell cycle exit

We began this study with the primary goal of identifying chromatin remodeler(s) that may be responsible for decommissioning chromatin at three rate-limiting cell cycle genes after cell cycle exit occurs. As we identified roles for Mi-2 and E93 in promoting cell cycle exit we examined accessibility at each of these loci under conditions where Mi-2 or E93 are disrupted. As mentioned above, we observed no significant effect of Mi-2^DN^ on chromatin accessibility at *cycE* and minor effects at *e2f1* (Figure S6). Although this ruled out a major role for Mi-2 in mediating the decommissioning that occurs at these loci, when examining the E93^MUT^ wing FAIRE-seq data, we observed several peaks that show differential accessibility at each of these loci (Figure S9), suggesting E93 may coordinate with other, yet-to-be-identified chromatin remodelers to modulate dynamic elements at *e2f1* and *cycE*.

While examining the *string* locus, we observed a number of E93-bound regions that exhibit differential accessibility in E93^MUT^ wings, including a region overlapping the uncharacterized element that we identified earlier in the Mi-2^DN^ accessibility data from the eye (Figure 3E-F; Figure 7A). Consistent with its dynamics in the eye, this region is most accessible at the 24h APF time point in the wing and has largely closed by 44h APF (Figure 7A). Synthesizing our findings at this region across datasets, we observe: developmental dynamics that are mirrored across wing and eye tissues; E93 binding at 24h APF in the wing; a failure of decommissioning for this region with Mi-2^DN^ in the eye; and a similar persistence of accessibility with loss of E93 in the wing (Figure 7B). These data are suggestive that this distal element at *string* could be a shared target of Mi-2 and E93. Therefore, we chose to characterize this region further and tested it for regulatory activity in a transgenic reporter assay. Importantly, the reporter was constructed using a nuclear-localized tdTomato protein that includes a PEST degradation sequence, to reduce the protein half-life and allow resolution to detect enhancer shut-off [53]. The reporter expression was assessed in eyes and wings; in both tissues it is inactive at larval stages but exhibits dynamic reporter activity during metamorphosis in the hours after cell cycle exit, with a peak of reporter expression at 30-36h followed by a decline by 48-54h APF (Figure 7C-D, Figure S10A-B). These dynamics are consistent with the decommissioning of this enhancer observed via chromatin accessibility assays from 24h to 44h in both tissues. Overall, these results support the identification of a novel **P**upal **E**nhancer element at the *stg* regulatory locus, which we refer to as *stg*-PE.

**Figure 7:**
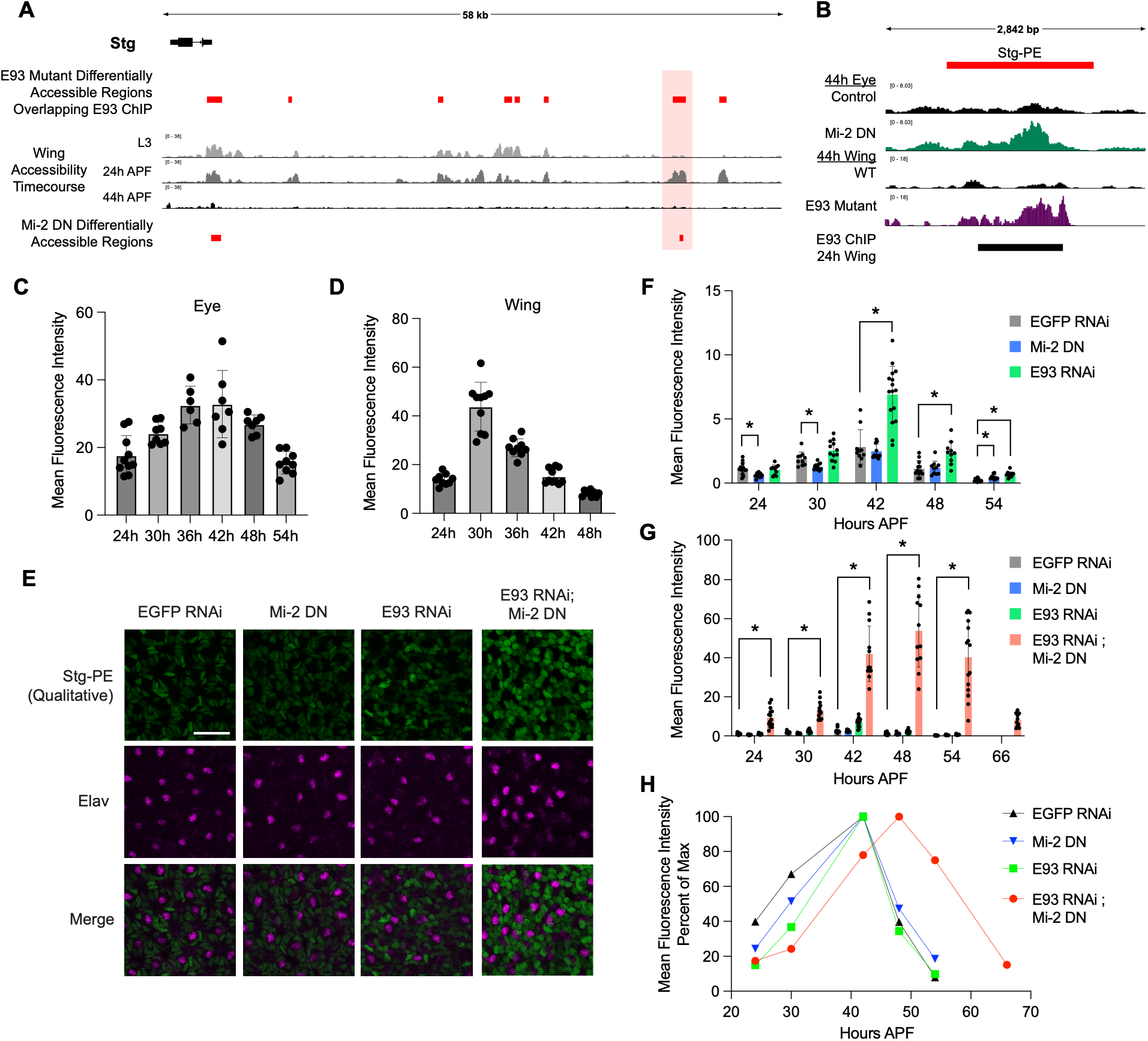
Mi-2 and E93 synergistically regulate Stg-PE, a novel *string* pupal enhancer. (**A**) Diagram of the *stg* genomic locus and upstream regulatory region. ATAC-Seq data from a time course of wings at L3, 24h and 44h APF. Differentially accessible regions in 44h APF *E93* mutant wings that overlap 24h APF E93 binding sites identified via ChIP-seq are indicated above by red boxes. Differentially accessible regions in 44h APF Mi-2 DN expressing eyes as from Figure 3E are indicated in red boxes below. (**B**) View of the region highlighted with the light red box in panel A, showing ATAC-Seq data from control and Mi-2^DN^-expressing eyes at 44h APF and FAIRE-Seq data from control and *E93* mutant wings at 44h APF. Y-axes indicate count-per-million normalized FAIRE-Seq or ATAC-Seq coverage. Black box below indicates region bound by E93 in 24h APF wing ChIP-Seq data. Red box above indicates the Stg-PE region cloned for enhancer reporter assays. (**C**-**D**) Quantifications of nls-tdTomato-PEST signal driven by Stg-PE in the pupal eye (C) and wing (D) at indicated hours APF. (**E**) Representative images of Stg-PE driven nls-tdTomato-PEST reporter (green; note that images show qualitative signal) and anti-Elav signal (magenta) in the basal layer of 42h APF eyes expressing EGFP RNAi, Mi-2 DN, E93 RNAi, or combined expression of E93 RNAi and Mi-2 DN. (**F**-**G**) Quantifications of Stg-PE reporter signal from images as in panel E at the indicated hours APF. EGFP RNAi, grey bars; Mi-2 DN, blue bars; E93 RNAi, green bars; E93 RNAi and Mi-2^DN^ combination, red bars. Each data point represents one eye; bar height indicates average and error bars depict standard deviation. Asterisks mark statistically significant comparisons between the EGFP RNAi control and each experimental condition: multiple unpaired t-tests with FDR correction, q<0.005. Note that the data in panel F are reproduced in panel G with the inclusion of the combined manipulation to aid in visual comparison between conditions. (**H**) Quantification of Stg-PE reporter signal as in panel G, scaled to the percent of maximum average signal in each condition. EGFP RNAi, black line; Mi-2 DN, blue line; E93 RNAi, green line; E93 RNAi and Mi-2^DN^ combination, red line.

To determine if Mi-2 and E93 regulate the enhancer activity of *stg*-PE in addition to modulating its accessibility, we next tested the reporter in the presence of Mi-2^DN^ or E93 RNAi in the eye. As the predominant *stg*-PE reporter signal in the eye comes from the interommatidial pigment cell nuclei in the base of the eye (Figure S10C), we performed our quantifications from images taken in this plane of the tissue (Figure 7E shows representative qualitative reporter signal). These quantifications revealed that both Mi-2^DN^ and E93 knockdown are sufficient to modulate the activity of *stg*-PE (Figure 7F). Reporter intensity at the 24h and 30h APF time points is reduced with Mi-2^DN^ relative to control. The reporter then reaches control levels at its peak and ultimately shows persistent activity at 54h APF. These results suggest that Mi-2 supports the initial activation of Stg-PE and also ensures this element’s complete shut-off. In contrast, E93 RNAi does not alter *stg*-PE activity at the early time points but induces a marked increase in enhancer activity at its peak followed by persistent residual reporter expression at each time point afterward. This suggests that E93 primarily acts to restrain *stg*-PE’s peak activity level. Finally, we wondered whether combining Mi-2^DN^ with E93 knockdown might further modulate the activity of *stg*-PE; indeed, combining these manipulations induces a striking de-repression of *stg*-PE’s enhancer activity at every time point and delays the peak of its activity to 48h APF (Figure 7G-H). We note that in control eyes the *stg*-PE reporter falls to less than 50% of maximum signal 6 hours after its peak, suggesting a relatively swift shut-off (Figure 7H). However, with the combined manipulation, the reporter retains 75% of maximum signal 6 hours after peak; this is consistent with the idea that in addition to dramatically derepressing *stg*-PE, combining Mi-2^DN^ with E93 knockdown also compromises this element’s decommissioning. As a whole, these data support the identification of a novel pupal enhancer at the *string* locus that is synergistically regulated by Mi-2 and E93, suggesting that these two factors together ensure the postmitotic decommissioning of this critical cell cycle regulator.

## Discussion

### Mi-2 and E93 alter chromatin accessibility and gene expression to coordinate cell cycle exit with progression of terminal differentiation

Our data support a model for how the general chromatin remodeler Mi-2 and the transcriptional cascade downstream of ecdysone signaling together promote stable cell cycle exit in coordination with the progression of terminal differentiation in wings and eyes (Figure 8). Mi-2 catalyzes the movement of nucleosomes at developmentally dynamic regions throughout the genome, primarily supporting the timely execution of chromatin decommissioning events. The ecdysone target E93 is upregulated in response to a high threshold of ecdysone signaling and works cooperatively with Mi-2 to co-regulate accessibility at a select number of shared target regions during the final cell cycle, including cell cycle and terminal differentiation genes. In particular, Mi-2 and E93 each support decommissioning events at the early differentiation gene *broad* and key mitotic regulator *string* to facilitate their downregulation, ensuring a robust cell state transition from early to late differentiation and from a proliferative to a stable postmitotic state. Moreover, we find that these factors synergistically regulate the activity and shut-off of a novel *string* enhancer that is specifically utilized in the pupal stages.

**Figure 8:**
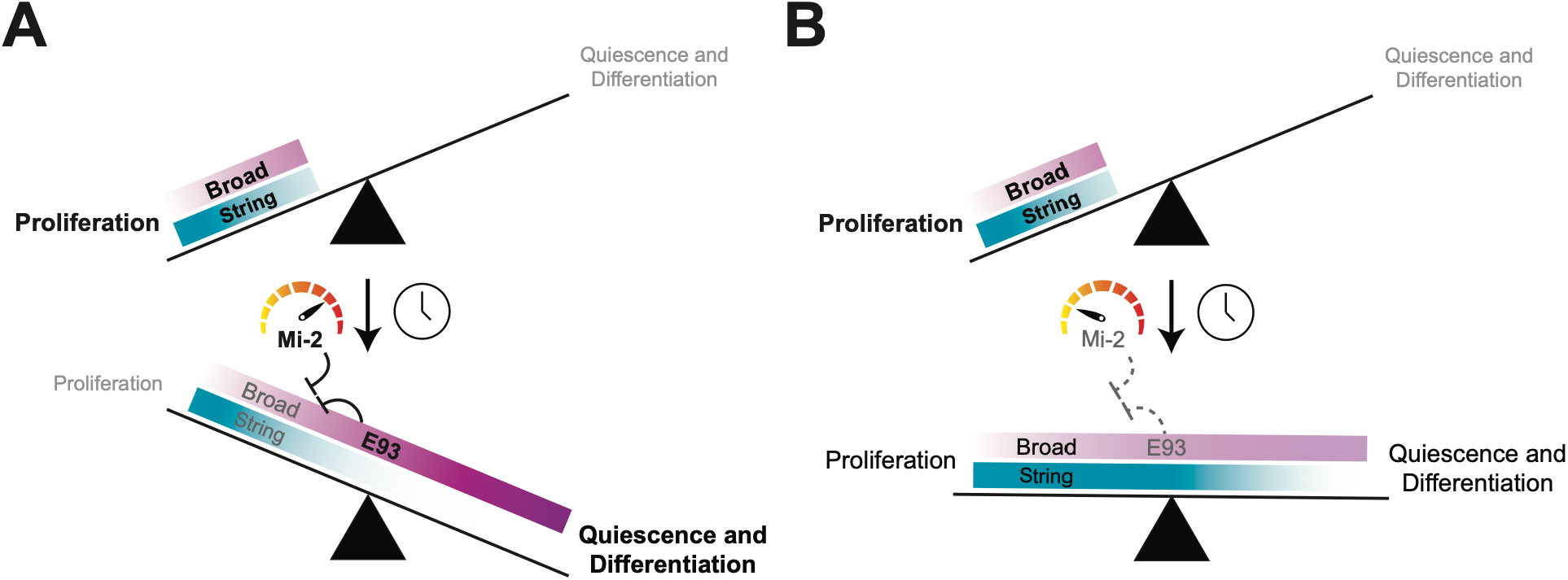
Mi-2 and E93 work cooperatively to coordinate termination of the cell cycle with initiation of differentiation. (**A**) Conceptual model depicting the cell state transition that occurs as proliferation ends and terminal differentiation proceeds. The magenta gradient represents the ecdysone transcriptional cascade while the blue gradient represents cell cycle gene expression. Mi-2 modulates the speed of developmental transitions and works with the temporal factor E93 to decommission the cell cycle gene *String* as well as the early differentiation factor *Broad*. (**B**) Manipulations that interfere with Mi-2 or E93 function compromise this transition, resulting in persistent cell cycle and early differentiation gene expression and muted progression into terminal differentiation.

Over 15 years ago, the existence of exit elements was postulated – regulatory elements that mediate the robust transcriptional shut down of cell cycle genes during cell cycle exit and terminal differentiation [9]. Exit elements could mediate transcriptional repression through binding repressors or through recruiting chromatin remodelers leading to a repressive chromatin state [55]. For most cell cycle genes, the regulatory elements involved in repression are also involved in gene activation. This is best resolved for targets of RB/E2F regulation where the same promoter-proximal element mediates E2F-dependent gene activation as well as RB or DREAM-mediated repression [30]. But the rate-limiting cell cycle genes with more complex cis regulatory modules could be subject to the control of exit elements to maintain repression during stable postmitotic states. Our previous work demonstrates that the stable repression of rate-limiting cell cycle genes likely involves enhancer decommissioning through loss of transcription factor binding and nucleosome movement, rather than formation of repressive chromatin via heterochromatin-mediated repression [56]. Importantly, we also showed that the decommissioning of several potential regulatory elements at rate-limiting cell cycle genes was developmentally controlled, and independent of the cycling status of the terminally differentiating cells [13]. This raised longstanding questions of how the decommissioning at rate-limiting cell cycle genes is coordinated with the terminal differentiation process and which effectors are responsible for the chromatin modulation. While the work here shows a role for the remodeler Mi-2 in the proper shut down of the mitotic regulator *string*, future work will be needed to identify the remodeler(s) responsible for decommissioning regulatory elements in the critical G1-S regulatory gene *cyclin E*.

Mi-2 remodels nucleosomes by moving end nucleosomes towards the center of a DNA fragment and is known to bind to nucleosomal DNA through its chromodomains [57, 58]. Mi-2 is also known to function as a member of two complexes in *Drosophila melanogaster*, the Nucleosome Remodeling and Deacetylase complex (NuRD) and the Drosophila MEP-1 complex (dMec). Both complexes are thought to serve mostly transcriptional repressive functions, with the NuRD complex having a histone deacetylase function while dMec uses a histone deacetylase-independent mechanism [33, 59]. In our RNAi screen for cell cycle exit regulators we tested HDAC1, the main HDAC in the NuRD complex, as well as Caf1-55 and MBD-like. None of these NuRD knockdowns resulted in any obvious cell cycle exit phenotype, despite robust accumulation of H3K27 acetylation with the HDAC1 knockdown. We therefore suggest that the functions of Mi-2 described here may be NuRD independent, although we cannot rule out the possibility that NuRD plays a role in maintaining cell cycle exit at timepoints later than those examined here.

### Disruption of E93/Mi-2 function leads to a block in the ecdysone feed-forward cascade

Inhibition of either E93 or Mi-2 led to a disruption of the normal ecdysone feed-forward transcriptional cascade in eyes and wings (this work and [6]). E93 expression is activated around 12h APF, around the time of transition from prepupal to pupal stages [60]. The E93 binding motif is enriched in chromatin accessibility peaks that close between third larval instar (L3) and 24h APF as well as peaks that open between 24h and 44h APF [6]. This suggests that E93 may play a role in not only activating gene expression at later developmental stages, but may aid in closing chromatin at genes expressed earlier in development. When expressed ectopically at earlier stages of development, E93 has also been shown to prematurely turn off early stage enhancers while binding to and opening late stage enhancers [61]. In the case of E93 loss, the early ecdysone targets Broad and Ftz-F1 remain high, presumably leading to subsequent temporal defects in the execution of the proper cascade. In the case of blocking Mi-2 function, Broad and Ftz-F1 also remain high, although with milder temporal cascade defects. Interestingly, when Mi-2 function is inhibited, E93 is upregulated (Fold Change 1.71) and we observe two significant sites of chromatin accessibility upstream of E93 that fail to close. This suggests E93 is upregulated in response to Mi-2 loss of function, which may compensate to correct the chromatin accessibility changes and gene expression changes through E93 acting in concert with other chromatin remodelers. Consistent with this, the wing and eye phenotypes and effects on cell cycle exit are much stronger for E93 inhibition than Mi-2 inhibition. There is abundant evidence of redundancies and chromatin remodelers acting in concert [62], which may be uncovered through additional manipulations during cell cycle exit.

Notably, prior work that has implicated Mi-2 in the expression of ecdysone targets in cell culture characterized its role as a general repressor of target transcripts in the absence and presence of hormone [35]. However, rather than a general upregulation of all ecdysone targets with the expression of Mi-2^DN^, we observe a more nuanced slowing or stalling of the cascade’s temporal progression. Based on our *in vivo* studies, we propose that Mi-2 modulates the ecdysone cascade’s temporal progression through its coregulatory action with E93.

## Acknowledgements

We thank Dr. Fernando Casares for generously providing reporters for many Stg regulatory elements. We thank Dr. John Tamkun for kindly providing the UAS-Mi2^WT^ and UAS-Mi2^DN^ lines. We thank Dr. Helena Richardson for kindly providing the GMR-Gal4, UAS-CycE; GMR-P35 line. We thank the UM Advanced Genomics Core for assistance with sample quality control, RNA-seq library preparation and sequencing. Additional fly stocks obtained from the Bloomington Drosophila Stock Center (NIH P40OD018537) were used in this study. We used FlyBase (release FB2024_03) to find information on phenotypes/stocks/gene expression. Antibodies were obtained from the Developmental Studies Hybridoma Bank, created by the NICHD of the NIH and maintained at the University of Iowa, Department of Biology, Iowa City, IA 52242. EMB was supported by the U. Michigan Genetics Training Grant (T32GM007544). This work in the Buttitta Lab was supported by the National Institute of General Medical Sciences of the National Institutes of Health (R35GM149273). Work in the McKay lab is supported by the National Institute of General Medical Sciences (R35GM128851).

## Methods

### Fly Stocks

Figure 1 genotypes:

1A and B: Sensitized: *w; GMR-Gal4, UAS-CycE/+; GMR-P35/UAS-GFP^RNAi^* (UAS-GFP^RNAi^ from BDSC 9330)

Suppressed: *w; GMR-Gal4, UAS-CycE/+; GMR-P35/UAS-Wee, UAS-Dap* (UAS-Wee+Dap described in [63])

Enhanced: *w; GMR-Gal4, UAS-CycE/+; GMR-P35/UAS-Mi-2^RNAi^* (Mi-2^RNAi^ BDSC 51774) and *w; GMR-Gal4, UAS-CycE/+; GMR-P35/Mi2^4^* (Mi2^4^ allele BDSC 26170)

CycD/Cdk4/ E2F: *w; GMR-Gal4, UAS-CycE/UAS-CycD, UAS-Cdk4; GMR-P35/UAS-E2F1, UAS-Dp* (UAS-CycD, UAS-Cdk4; UAS-E2F1, UAS-Dp described in [13]).

1C: *w; en-Gal4, UAS-GFP/UAS-E2F1, UAS-DP; tub-Gal80^TS^/UAS-white^RNAi^ or Mi-2^RNAi^* (white^RNAi^ BDSC BL35573). Animals were shifted from 18 deg to 28deg at early L3. Wings are 28h APF (corrected for relative timing at 25°C as described in [9]).

1D-F: *w; GMR-Gal4, UAS-CycE/+; GMR-P35/+ or GMR-Gal4, UAS-CycE/+; GMR-P35/UAS Mi-2^WT^ or Mi-2^DN^* (Mi-2 lines kindly provided by J. Tamkun, described in [37])

1G *w; en-Gal4, UAS-GFP; tub-Gal80^TS^/UAS-Mi-2^DN^* Animals were shifted as described in 1C.

ATAC-seq and RNA-seq genotypes:

*GMR-P35/w^1118^; GMR-Gal4/+; +/+*

*GMR-P35/+; GMR-Gal4/+; UAS-Mi-2^WT^-FLAG/+*

*GMR-P35/+; GMR-Gal4/+; UAS-Mi-2^DN^-FLAG/+*

*GMR-P35/+; GMR-Gal4/+; UAS-Mi-2^RNAi^/+*

*GMR-P35/+; GMR-Gal4/+; UAS-mCherry^RNAi^/+* (mCherry^RNAi^ BDSC BL35785)

*GMR-P35/+; GMR-Gal4/UAS-E93^RNAi^/+; +* (E93^RNAi^ Vienna *Drosophila* Resource Center, #104390)

All crosses were kept at 25°C.

Figure 6 genotypes:

6A: *GMR-P35/w^1118^; GMR-Gal4/+; +/+*

*GMR-P35/+; GMR-Gal4/UAS-E93^RNAi^/+; +*

6B-C: *w^1118^/+; GMR-Gal4, UAS-CycE/+;GMR-P35/+*

*+; GMR-Gal4,UAS-CycE/UAS-E93^RNAi^; GMR-P35/+*

All crosses were kept at 25°C.

Figure 7 genotypes:

7C-D: *w; +; StgPE-tdTomatoPEST*

7E-H: *GMR-P35; GMR-Gal4/UAS-EGFP^RNAi^; StgPE-tdTomatoPEST/+* (EGFP^RNAi^ BDSC 41555)

*GMR-P35; GMR-Gal4/+; StgPE-tdTomatoPEST/UAS-Mi-2^DN^-FLAG*

*GMR-P35; GMR-Gal4/UAS-E93^RNAi^; StgPE-tdTomatoPEST/+*

*GMR-P35; GMR-Gal4/UAS-E93^RNAi^; StgPE-tdTomatoPEST/UAS-Mi-2^DN^-FLAG*

Supplemental Figure 1 genotypes:

S1A: *w; GMR-Gal4/+; UAS-white^RNAi^* or *UAS-Mi-2^DN^/ PCNA-GFP* (white^RNAi^ BDSC BL35573, PCNA-GFP from [30])

S1B: *w; en-Gal4, UAS-GFP; tub-Gal80TS/UAS-white^RNAi^*or *Mi-2^RNAi^*. Animals were shifted from 18°C to 28°C at early L3. Wings are 28h APF (corrected for relative timing at 25°C as described in [9]).

S1C: *w; GMR-Gal4, UAS-CycE/+; GMR-P35/+* or *GMR-Gal4, UAS-CycE/+; GMR-P35/UAS-Mi-2^WT^* and genotypes described in Fig. 2 A.

S1D: *w^1118^;GMR-Gal4/+;+* or *w^1118^;GMR-Gal4/+; UAS-Mi2^WT^ or ^DN^* as described for Figure 1.

Supplemental Figure 8 genotypes:

S8A: *w1118;+;E93^mimicGFP^*(E93^mimicGFP^ BDSC BL43675)

S8B: *w^1118^; en-Gal4/UAS-E93^RNAi^; tub-Gal80^TS^/E93^mimicGFP^* Wings are 26h APF (eggs were laid and raised at 18°C, WPP were collected and raised at 29°C until an equivalent of 26h APF)

S8C: *GMR-P35/+; GMR-Gal4/UAS-E93^RNAi^; +* or *GMR-P35/+; GMR-Gal4/+ UAS-mCherry^RNAi^/+*

S8D: *w^1118^/GMR-P35; GMR-Gal4/UAS-E93^RNAi^; +*

Supplemental Figure 10 genotypes:

S10A-B: *w; +; StgPE-tdTomatoPEST*

S10C: *GMR-P35; GMR-Gal4/UAS-EGFP^RNAi^; StgPE-tdTomatoPEST/+*

Generation of StgPE-tdTomatoPEST transgenic reporter lines:

The 1,611 bp StgPE amplicon (dm6 coordinates chr3R:29,299,505-29,301,115) was PCR amplified and Gateway cloned (Invitrogen) first into a pENTR vector and verified by restriction digest and Sanger sequencing. The region was then cloned into the pDEST-attR1/2-tdTomato-PEST vector [53] and verified by restriction digest and plasmid sequencing. The transgenic construct was inserted into the genome by PhiC31 integration at the attP40 and attP2 landing sites with injections performed by BestGene. Similar developmental dynamics for the reporter were observed regardless of insertion site.

### Immunohistochemistry and imaging

For pupal tissues, white prepupae were collected and raised at 25°C until the appropriate age for dissection unless otherwise noted above. Tissues were dissected in 1X PBS and fixed for 30 min in 1X PBS + 4% paraformaldehyde. S2 cells were plated on concavalin-A coated coverslips, fixed with 4% paraformaldehyde in 1X PBS. Primary antibodies were diluted in PAT (1X PBS containing 0.1% Triton X-100 + 1% BSA) and incubated for 4 hours at room temperature or overnight at 4°C. Secondary antibodies were diluted in PBT-X (1X PBS containing 0.3% Triton X-100 with 0.1% BSA) + 2% normal goat serum and incubated for 4 hours at room temperature or overnight at 4°C. DAPI was diluted 1:1000 in 1X PBS + 0.1% Triton X-100, then incubated with tissues for 10 minutes at room temperature in the dark. Samples were mounted onto glass slides using Vectashield mounting medium (Vector Laboratories). Slides were imaged on a Leica SP5, SP8 or Stellaris confocal microscope. Images were quantified in FIJI. PH3 was quantified using max projections of z stacks, and using a minimum signal intensity threshold of 20 to count puncta. Average tdTomato reporter intensity was measured from max projections of z stacks to assess expression of Stg-PE reporter in Figure 7C-D, while quantifications were done from single z slices for Figure 7F-H.

### Antibodies

Mouse Anti-Discs large (4F3), DSHB, 1:100

Rabbit Anti-Phosphohistone H3 (serine 10) (D2C8), Cell Signaling, cat#3377, 1:1000

Rat Anti-Elav (7E8A10), DSHB, 1:100

Rabbit Anti-GFP, Invitrogen, cat#A11122, 1:1000

Mouse Anti-HA, Sigma, clone HA-7, 1:2000

Goat Anti-Mouse Plus (488), AlexaFluor, cat#A32723, 1:2000

Goat Anti-Rabbit (488), AlexaFluor, cat#A11034, 1:2000

Goat Anti-Rabbit (568), AlexaFluor, cat#A11036, 1:2000

Goat Anti-Rat (568), AlexaFluor, cat#A11077, 1:2000

AlexaFluor 647 conjugated streptavidin, Invitrogen, cat#S32357, 1:1000

### ATAC-seq & RNA-seq

For both ATAC-seq and RNA-seq, crosses were kept at 25°C. White prepupa were collected, sexed, and aged at 25°C until dissection at either 24h or 44h APF. ATAC-seq was performed on 24h and 44h APF eyes using pupal tissue dissociation and ATAC-seq protocol as described in [64]. Approximately 16 eyes were used per sample, with 2-3 replicates per genotype and timepoint.

For RNA-seq, 16 eyes were used per sample, with three replicates for GMR x w^1118^, Mi-2^WT^ and Mi-2^DN^ genotypes 24h and 44h APF timepoints, and four replicates for GMR x mCherry^RNAi^ and E93^RNAi^ 44h APF samples. Pupal eyes were dissected in filtered 1X PBS, then transferred to 400 µL Trizol containing 0.5 µL of 20mg/mL glycogen.

Samples were vortexed for 2 minutes, then 80 µL of chloroform was added and samples were vortexed, then centrifuged for 15 minutes at 12,000xg at 4°C. Aqueous layer was transferred to a new Eppendorf tube, then 250 µL of Isopropanol was added. Samples were mixed, then frozen overnight at −20°C. The next day, samples were centrifuged for 5 minutes at 12,000xg at 4°C. Alcohol was removed and samples were resuspended in 50 µL of RNAse free water. Libraries were generated by the University of Michigan Sequencing Core. Total RNA was sequenced for GMR x w^1118^, Mi-2^WT^ and Mi-2^DN^ samples, and polyA selection was performed for GMR x mCherry^RNAi^ and E93^RNAi^ samples. All sequencing was performed at the University of Michigan Sequencing Core. An Agilent Tape Station was used to assess library quality. All ATAC-seq and RNA-seq samples were sequenced using Illumina NovaSeq S4 300 cycles, paired-end 150bp reads with a target depth of 90-100 million reads. ATAC-seq and RNA-seq data was analyzed as described in [64] with the following changes: Statistics were performed and ATAC-seq multidimensional scaling plots were generated using R package edgeR [65]. RNA-seq reads were mapped to the dm6 genome using STAR 2.7.6a [66]. Line plots and heatmaps of accessibility data were generated with deeptools utilities [67]. Genomic datasets were processed and intersected using bedtools utilities [68]. Randomization-based permutation test for statistical analysis of overlap between datasets was performed using regioneR [69].

### E93 TurboID

Three different inducible cell lines were generated using the pMT-puro vector: TurboID alone, E93 alone, E93+TurboID. Cells were cultured in biotin-depleted Schneider’s media containing 2ug/mL puromycin. Protein expression was induced with CuSO4 for 16-hours. Cells were washed in 1XPBS and incubated in media containing 500uM exogenous biotin for 2-hours. Biotin-minus media was used as an additional control for E93+TurboID cells. Nucleoplasmic and chromatin-bound proteins were extracted (Thermo, cat#78840). Biotinylated proteins were pulled down with magnetic streptavidin beads (Thermo, cat#65601) overnight at 4C. Washed beads were submitted to the UNC Proteomics Core Facility for mass spectrometry. Samples were prepared via on-bead trypsin digestion, followed by LC-MS/MS using Thermo Easy nLC 1200-QExactive HF.

### Data Access

The data generated in this study can be accessed from NCBI GEO, datasets GSE259302 (ATAC-Seq) and GSE259303 (RNA-Seq).

Previously published 24h and 44h APF wing E93 mutant FAIRE-seq and 24h APF wing E93^mimicGFP^ ChIP-seq data are available at GSE97956.

## Supplement to

**Figure S1:**
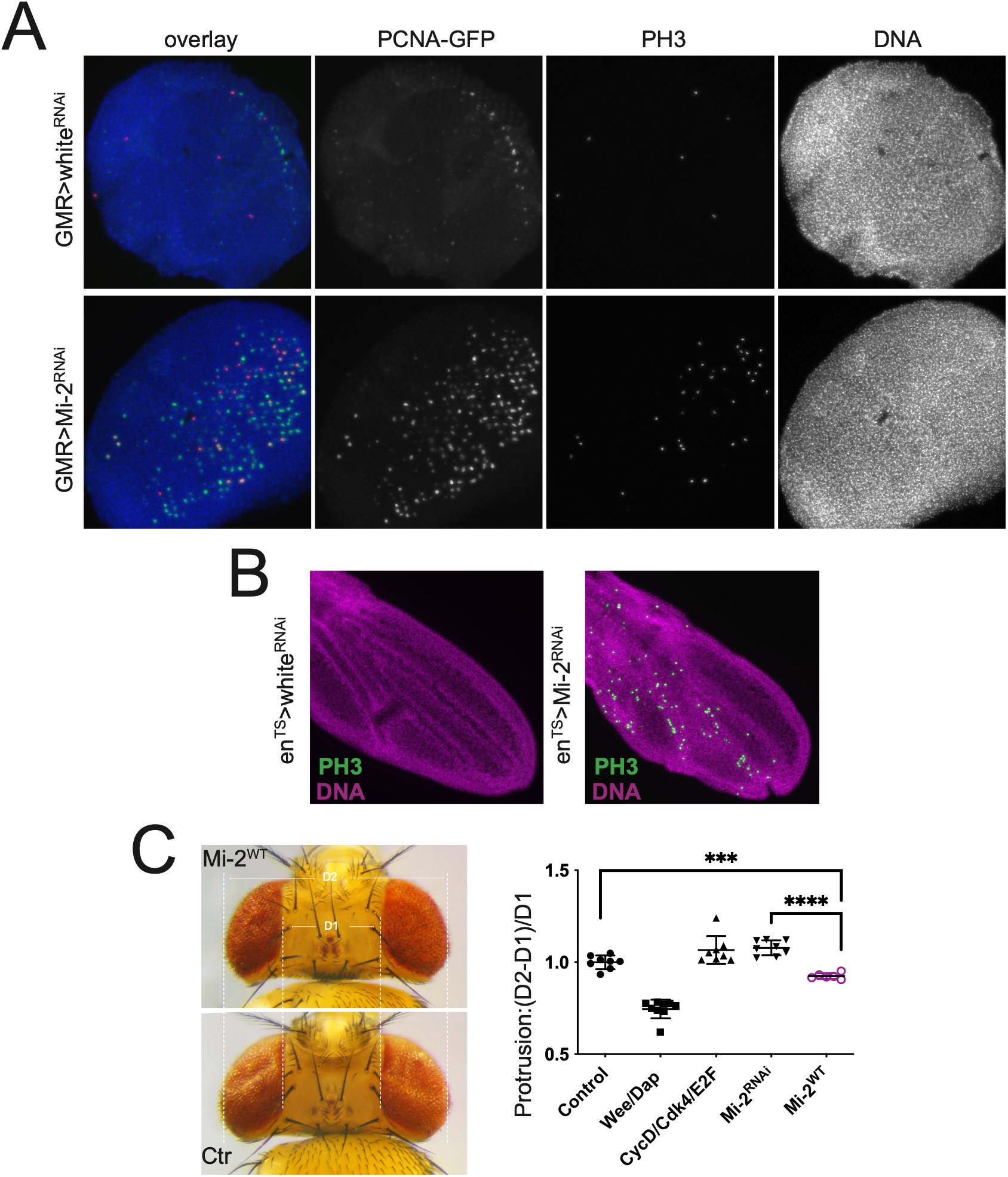
Additional evidence of Mi-2^RNAi^ and Mi-2^DN^ effects on cell cycle exit in the eye and wing. (**A**) Pupal eyes at 24h APF expressing a control (*white^RNAi^*) or *Mi-2*^RNAi^ during the final cell cycle under the control of *GMR*-Gal4 during the final cell cycle. DAPI marks DNA, PCNA-GFP reporter (Thacker and Duronio 2003) reads out E2F transcriptional activity and antibody recognizing phosphorylated Histone H3 (PH3) marks mitotic events. (**B**) 28h APF pupal wings expressing white^RNAi^ or Mi-2^RNAi^ in the posterior pupal wing using *en^TS^*. DAPI marks DNA (magenta) and antibody recognizing PH3 marks mitotic events (green). (**C**) Representative images (left) of adult eyes and eye protrusion measurements relative to control (right). Control (Ctr) sensitized eyes express GMR-P35 and CyclinE under the control of *GMR*-Gal4. Expression of suppressors and enhancers was performed in the sensitized background. Asterisks mark statistically significant comparisons; p-values derived from pairwise two-tailed t-tests.

**Figure S2:**
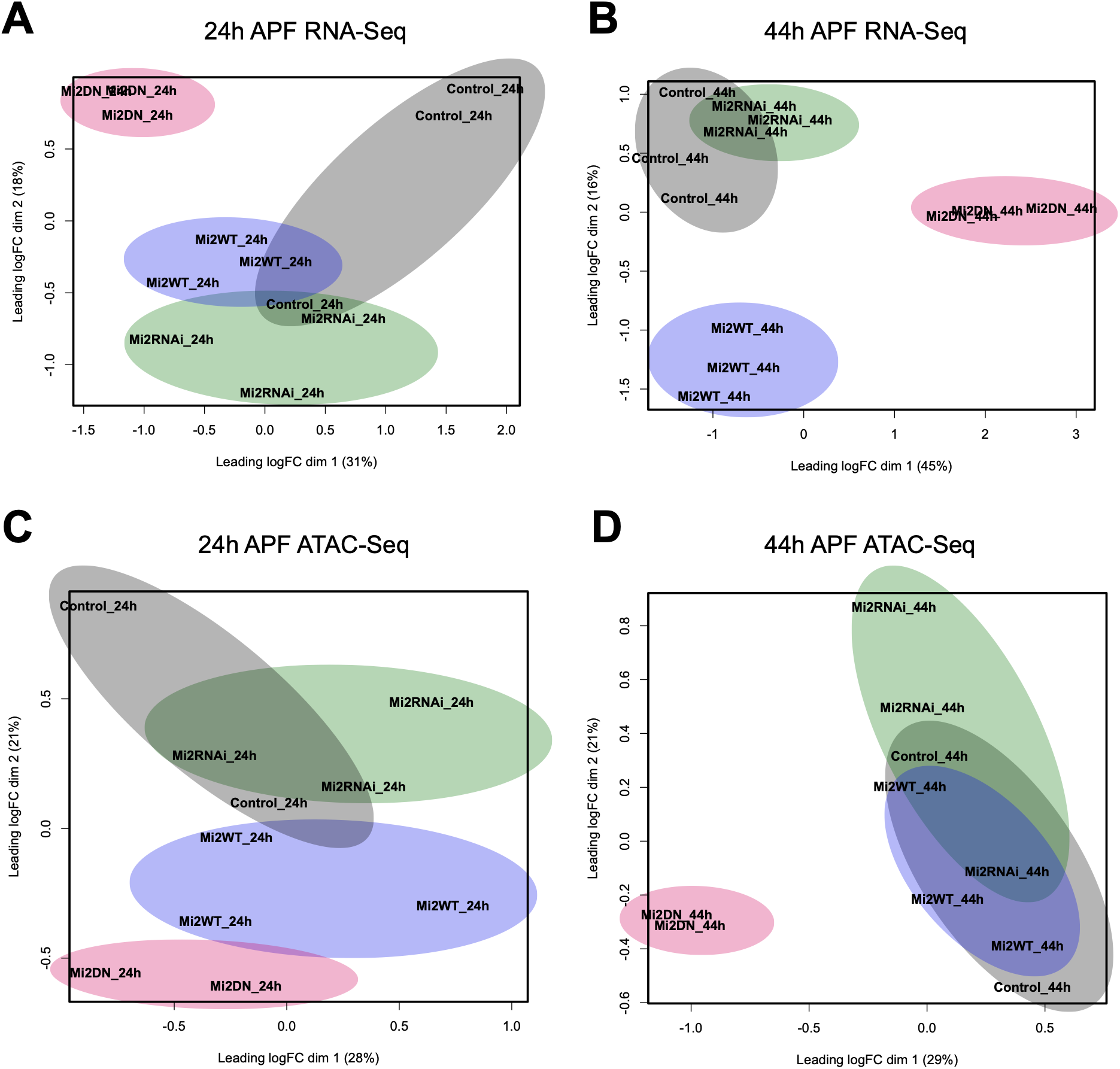
RNA-seq and ATAC-seq Multidimensional Scaling Plots. Multidimensional scaling plots for 24h APF (**A**) and 44h APF (**B**) RNA-Seq samples and 24h APF (**C**) and 44h APF (**D**) ATAC-Seq samples. Colored ovals highlight the clustering of each genotype relative to others.

**Figure S3:**
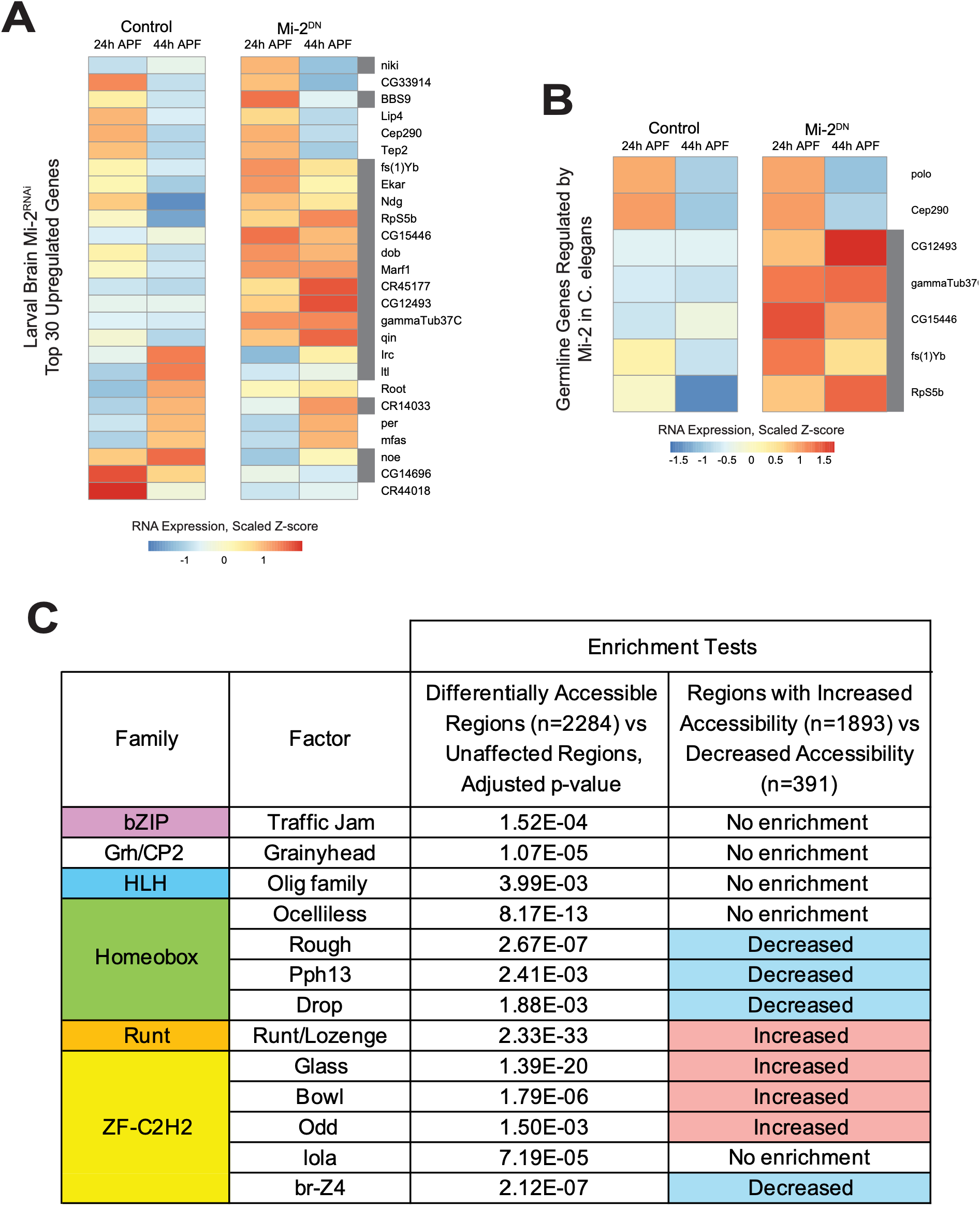
Mi-2^DN^ expression in the pupal eye recapitulates previously described functions for Mi-2 in other systems. (**A**) Gene expression heatmap showing the top upregulated genes following Mi-2 RNAi knockdown in larval brain as reported by Aughey et al, 2023. The expression of these genes as measured by RNA-Seq in the Control and Mi-2^DN^-expressing pupa eyes is plotted as scaled Z-score. Genes that were differentially expressed at 24h or 44h APF in Mi-2^DN^ relative to control are indicated by a grey box to the left of the gene name. (**B**) Gene expression heatmap showing germline genes reported to be repressed in somatic tissues by the homolog of Mi-2 in C. elegans in Unhavaithaya et al., 2002. The expression of these genes as measured by RNA-Seq in the Control and Mi-2^DN^-expressing pupa eyes is plotted as scaled Z-score. Genes that were differentially expressed at 24h or 44h APF in Mi-2^DN^ relative to control are indicated by a grey box to the left of the gene name. (**C**) Table highlighting select transcription factor binding motifs that are enriched among regions that exhibit differential accessibility with Mi-2^DN^ expression at 44h APF. All the motifs listed were enriched in a comparison of regions that show differential accessibility with Mi-2 DN compared to nonresponsive regions; some are differentially enriched when comparing regions with increased and decreased accessibility against one another. The factors highlighted here were selected based on previous literature describing a functional role in the developing eye; full datasets in Tables S1-S3.

**Figure S4:**
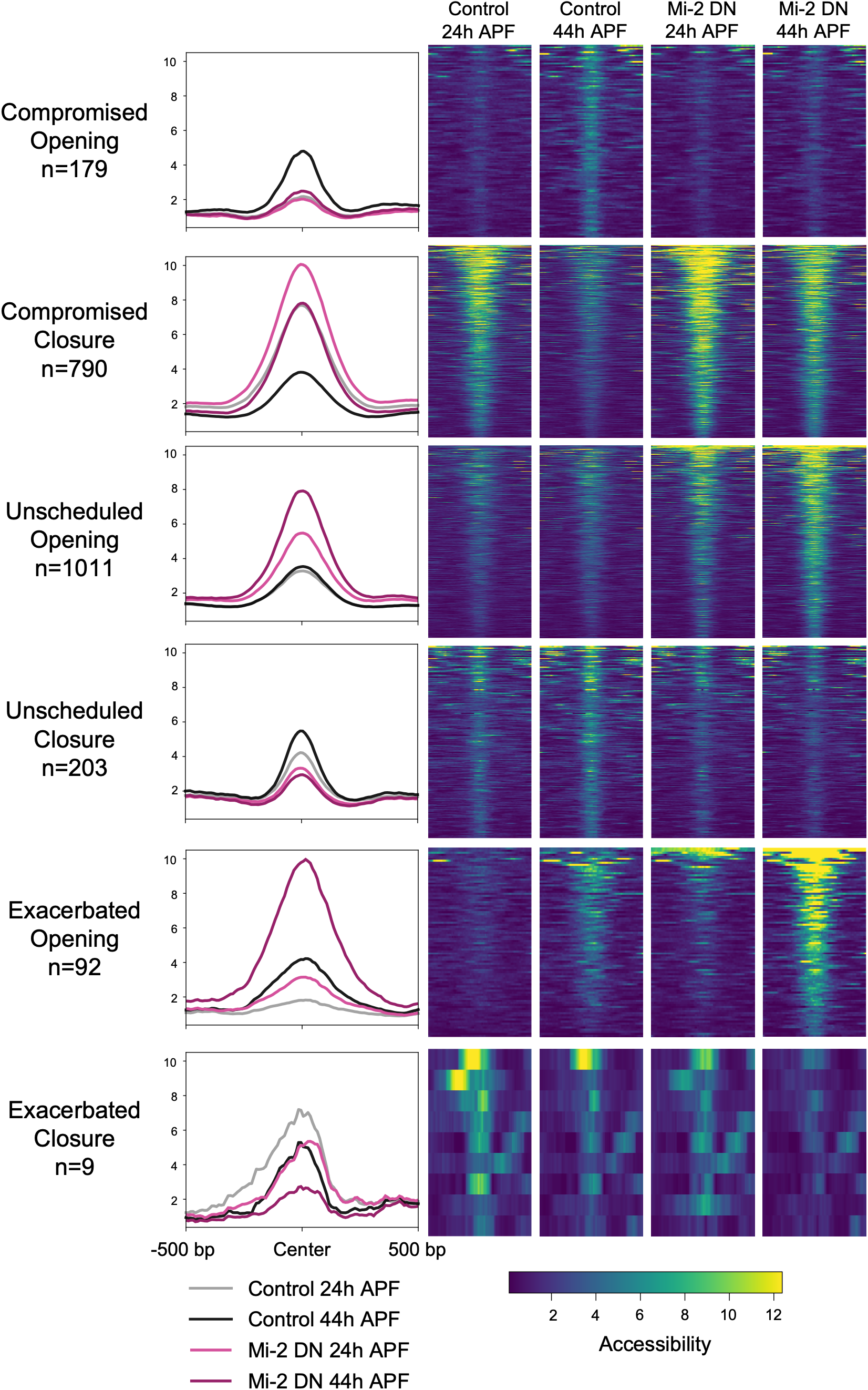
Accessibility changes for all Mi-2^DN^ dynamics profiles. Line plots and heatmaps showing the accessibility measures for regions falling into each of the dynamics profiles as indicated in Figure 2B. Line plots include samples at 24h (control: grey, Mi-2 DN: light pink) and 44h (control: black, Mi-2 DN: dark pink) and accessibility was averaged in 10 base pair bins over a 1 kilobase region centered on the peak center.

**Figure S5:**
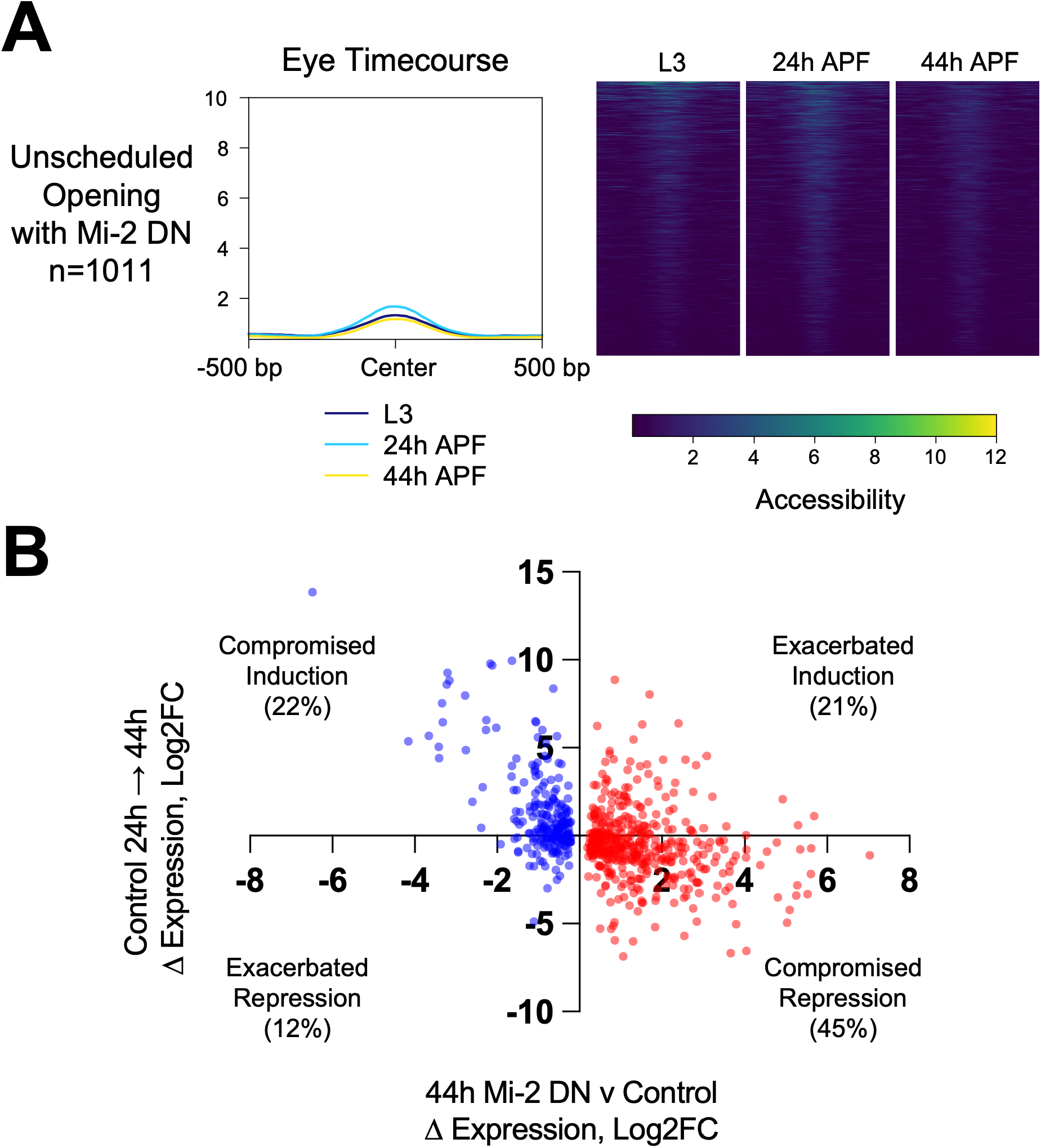
Accessibility and gene expression dynamics with Mi-2^DN^. (**A**) Line plot and heatmaps showing the accessibility measures in control time course samples for regions that exhibited unscheduled opening with Mi-2^DN^. Line plot shows averaged accessibility and includes samples at L3 (dark blue), 24h (light blue) and 44h (yellow) and accessibility was averaged in 10 base pair bins over a 1 kilobase region centered on the peak center. (**B**) Scatterplot showing the relationship between the change in expression (log2 fold change) induced by Mi-2^DN^ and the change in expression for the same genes in the control eye time course from 24h to 44h APF. Points are colored red (increased expression with Mi-2 DN) or blue (decreased expression with Mi-2 DN).

**Figure S6:**
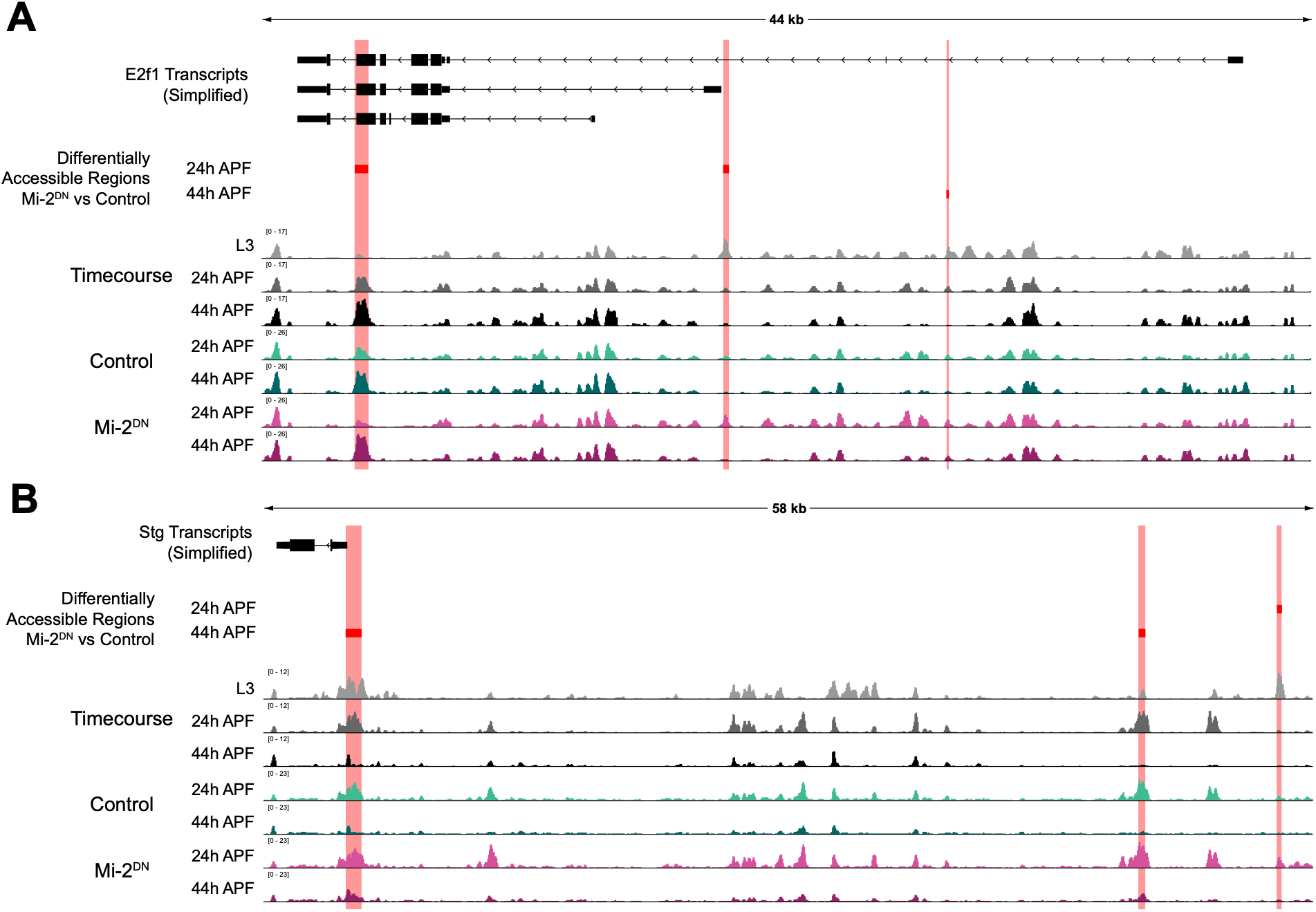
Mi-2^DN^ induces accessibility changes at E2f1 and String loci. Diagrams of the gene loci encoding *E2f1* (**A**) and *Stg* (**B**) with simplified RefSeq transcripts indicated in black. Data tracks depict ATAC-Seq data from a time course study of eyes at L3, 24h and 44h APF, as well as control and Mi-2^DN^ expressing eyes at 24h and 44h APF. Y-axes indicate count-per-million normalized read coverage. Regions that exhibit differential accessibility in Mi-2^DN^ relative to control at 24h or 44h APF are indicated with red boxes.

**Figure S7:**
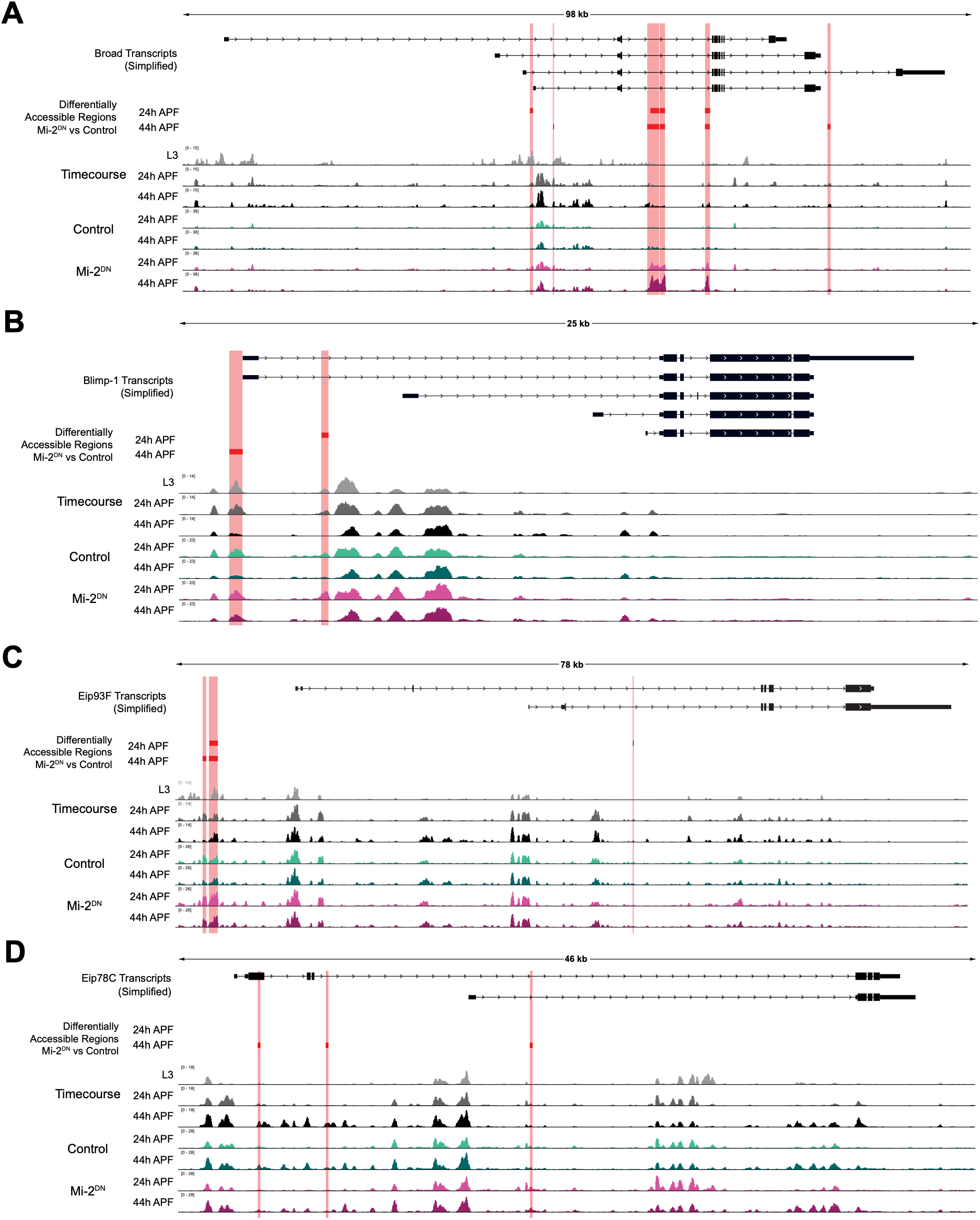
Mi-2^DN^ induces accessibility changes at ecdysone response loci. Diagrams of the gene loci encoding *Broad* (**A**), *Blimp*-1 (**B**), *Eip93F* (**C**) and *Eip78C* (**D**) with simplified RefSeq transcripts indicated in black. Data tracks depict ATAC-Seq data from a time course study of eyes at L3, 24h and 44h APF, as well as control and Mi-2^DN^ expressing eyes at 24h and 44h APF. Y-axes indicate count-per-million normalized read coverage. Regions that exhibit differential accessibility in Mi-2^DN^ relative to control at 24h or 44h APF are indicated with red boxes.

**Figure S8:**
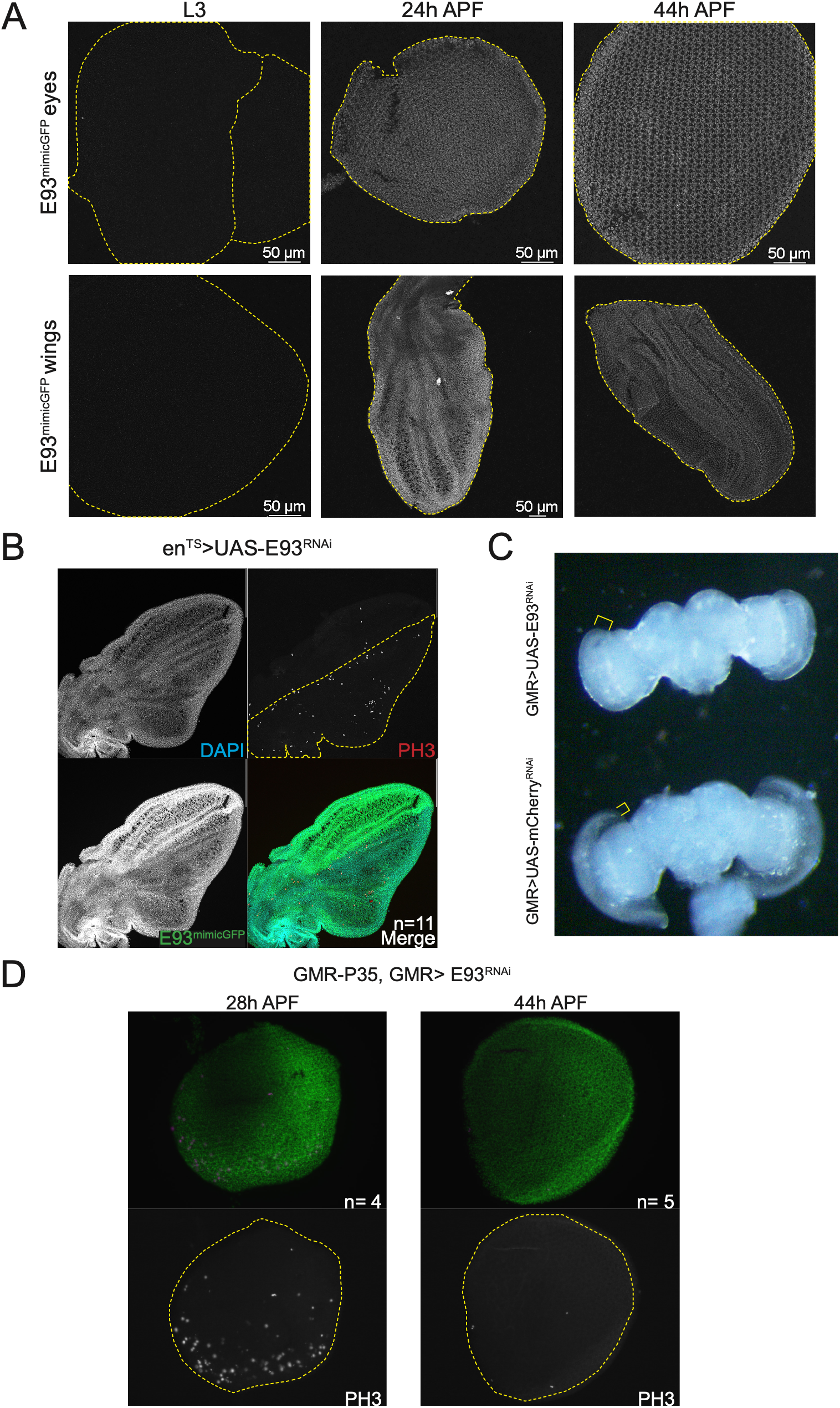
Characterization of E93 expression and E93^RNAi^-expressing eyes and wings. (**A**) Qualitative imaging of an endogenously tagged *E93* allele (E93^mimicGFP^) in larval and pupal eyes and wings. (**B**) 26h APF-equivalent wings expressing E93^RNAi^ in the posterior domain of the wing under the control of *en*-Gal4, *tub*-gal80^TS^ (en^TS^). DAPI marks DNA, an antibody against phosphorylated Histone H3 (PH3) marks mitotic events, and an antibody against GFP was used to amplify E93^mimicGFP^ signal. (**C**) Eye-brain complex at 45h APF from pupae expressing E93^RNAi^ or mCherry^RNAi^ under control of GMR-Gal4. Yellow brackets highlight the thickness of the developing retina in each condition. (**D**) 28h and 44h APF eyes expressing E93^RNAi^ under the control of GMR-Gal4. DAPI marks DNA (green) and antibody against PH3 marks mitotic events (grayscale).

**Figure S9:**
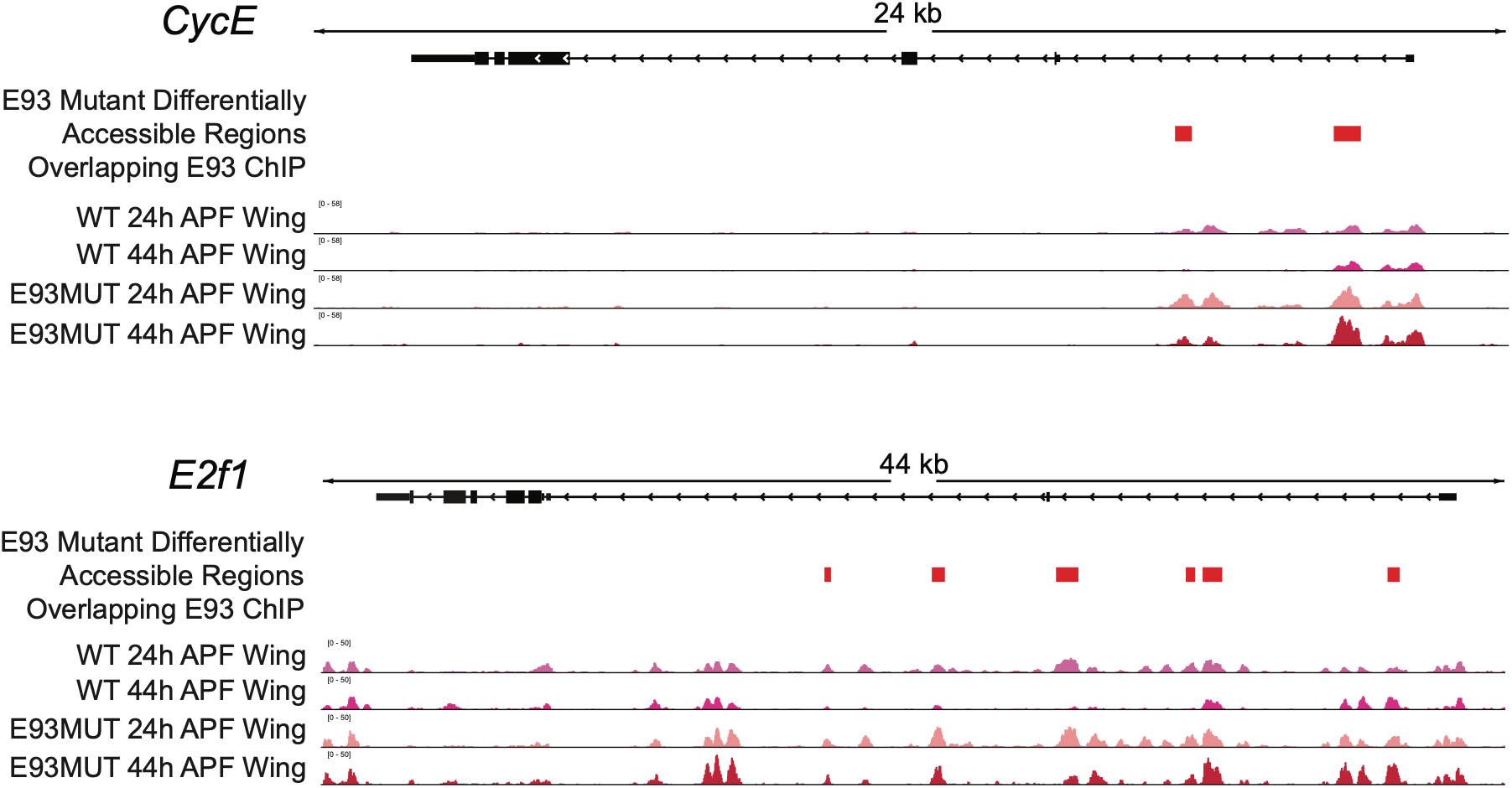
E93 mutants show accessibility changes at *e2f1* and *cycE.* Diagrams of the gene loci encoding *CycE* (**A**) and *E2f1* (**B**) with simplified RefSeq transcripts indicated in black. Data tracks depict FAIRE-Seq data from wildtype and E93 mutant wings at 24h and 44h APF. Y-axes indicate count-per-million normalized read coverage. Regions that exhibit differential accessibility in E93 mutant relative to control at 44h APF and that overlap E93 ChIP-Seq peaks are indicated with red boxes.

**Figure S10:**
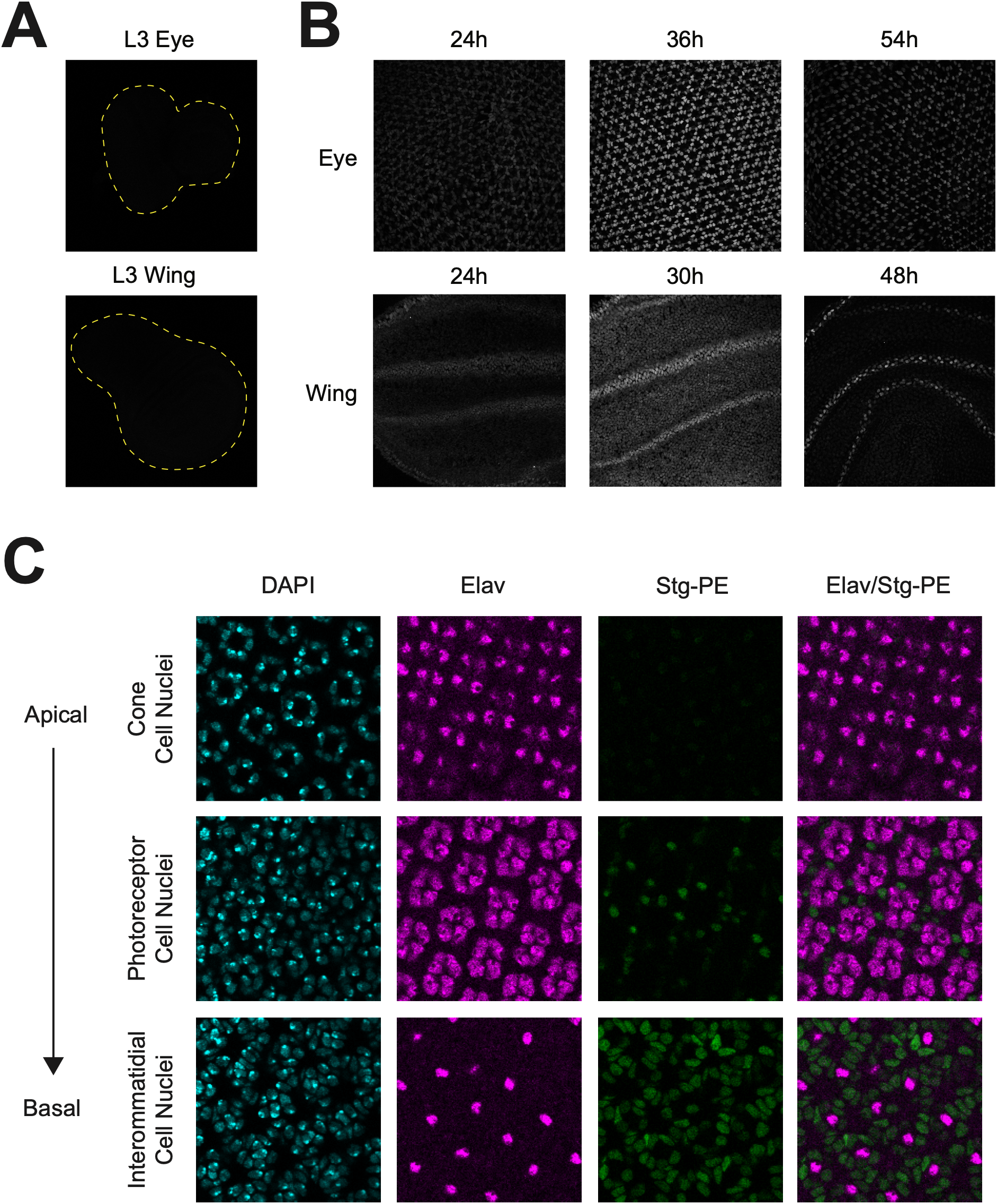
Stg-PE nls-tdTomato-PEST reporter expression in eyes and wings. (**A**) Eye-antenna disc and wing disc from late third instar larvae (outlined based on DAPI signal) show no tdTomato reporter expression. (**B**) Representative images of pupal eyes and wings showing dynamic tdTomato signal in the hours following cell cycle exit at the indicated hours APF. (**C**) Representative confocal images of control pupal eye at 42h APF imaged for DNA (DAPI), neurons (anti-Elav) and tdTomato reporter showing expression distribution from apical to basal layers of nuclei and from distinct cell types.

## Notes

### Competing Interest Statement

The authors have declared no competing interest.

## References

1. Niwa YS, Niwa R. Transcriptional regulation of insect steroid hormone biosynthesis and its role in controlling timing of molting and metamorphosis. Dev Growth Differ. 2016;58(1):94–105. Epub 20151215. doi: 10.1111/dgd.12248. PubMed PMID: 26667894; PubMed Central PMCID: PMCPMC11520982.

2. Thummel CS. Ecdysone-regulated puff genes 2000. Insect Biochem Mol Biol. 2002;32(2):113–20. Epub 2002/01/05. PubMed PMID: 11755052.

3. Thummel CS. Puffs and gene regulation--molecular insights into the Drosophila ecdysone regulatory hierarchy. Bioessays. 1990;12(12):561–8. doi: 10.1002/bies.950121202. PubMed PMID: 2127884.

4. Guo Y, Flegel K, Kumar J, McKay DJ, Buttitta LA. Ecdysone signaling induces two phases of cell cycle exit in Drosophila cells. Biol Open. 2016;5(11):1648–61. doi: 10.1242/bio.017525. PubMed PMID: 27737823; PubMed Central PMCID: PMCPMC5155522.

5. Uyehara CM, McKay DJ. Direct and widespread role for the nuclear receptor EcR in mediating the response to ecdysone in Drosophila. Proc Natl Acad Sci U S A. 2019;116(20):9893–902. Epub 20190424. doi: 10.1073/pnas.1900343116. PubMed PMID: 31019084; PubMed Central PMCID: PMCPMC6525475.

6. Uyehara CM, Nystrom SL, Niederhuber MJ, Leatham-Jensen M, Ma Y, Buttitta LA, et al. Hormone-dependent control of developmental timing through regulation of chromatin accessibility. Genes Dev. 2017. doi: 10.1101/gad.298182.117. PubMed PMID: 28536147.

7. Ninov N, Manjon C, Martin-Blanco E. Dynamic control of cell cycle and growth coupling by ecdysone, EGFR, and PI3K signaling in Drosophila histoblasts. PLoS Biol. 2009;7(4):e1000079. Epub 2009/04/10. doi: 08-PLBI-RA-2084 [pii] 10.1371/journal.pbio.1000079. PubMed PMID: 19355788; PubMed Central PMCID: PMC2672598.

8. Verma P, Cohen SM. miR-965 controls cell proliferation and migration during tissue morphogenesis in the Drosophila abdomen. Elife. 2015;4. doi: 10.7554/eLife.07389. PubMed PMID: 26226636; PubMed Central PMCID: PMCPMC4538364.

9. Buttitta LA, Katzaroff AJ, Perez CL, de la Cruz A, Edgar BA. A double-assurance mechanism controls cell cycle exit upon terminal differentiation in Drosophila. Dev Cell. 2007;12(4):631–43. doi: 10.1016/j.devcel.2007.02.020. PubMed PMID: 17419999.

10. Graves BJ, Schubiger G. Cell cycle changes during growth and differentiation of imaginal leg discs in Drosophila melanogaster. Dev Biol. 1982;93(1):104–10. PubMed PMID: 6813162.

11. Schubiger M, Palka J. Changing spatial patterns of DNA replication in the developing wing of Drosophila. Dev Biol. 1987;123(1):145–53. PubMed PMID: 3622926.

12. Milan M, Campuzano S, Garcia-Bellido A. Cell cycling and patterned cell proliferation in the Drosophila wing during metamorphosis. Proc Natl Acad Sci U S A. 1996;93(21):11687–92. PubMed PMID: 8876197.

13. Ma Y, McKay DJ, Buttitta L. Changes in chromatin accessibility ensure robust cell cycle exit in terminally differentiated cells. PLoS Biol. 2019;17(9):e3000378. Epub 2019/09/04. doi: 10.1371/journal.pbio.3000378. PubMed PMID: 31479438; PubMed Central PMCID: PMCPMC6743789.

14. Fogarty EA, Buchert EM, Ma Y, Nicely AB, Buttitta LA. Transcriptional repression and enhancer decommissioning silence cell cycle genes in postmitotic tissues. G3 (Bethesda). 2024;14(10). doi: 10.1093/g3journal/jkae203. PubMed PMID: 39171889; PubMed Central PMCID: PMCPMC11457063.

15. Kunert N, Brehm A. Novel Mi-2 related ATP-dependent chromatin remodelers. Epigenetics. 2009;4(4):209–11. Epub 20090506. doi: 10.4161/epi.8933. PubMed PMID: 19535903.

16. Albini S, Coutinho Toto P, Dall’Agnese A, Malecova B, Cenciarelli C, Felsani A, et al. Brahma is required for cell cycle arrest and late muscle gene expression during skeletal myogenesis. EMBO Rep. 2015;16(8):1037–50. Epub 2015/07/03. doi: 10.15252/embr.201540159. PubMed PMID: 26136374; PubMed Central PMCID: PMCPMC4552495.

17. Ruijtenberg S, van den Heuvel S. Coordinating cell proliferation and differentiation: Antagonism between cell cycle regulators and cell type-specific gene expression. Cell Cycle. 2016;15(2):196–212. doi: 10.1080/15384101.2015.1120925. PubMed PMID: 26825227; PubMed Central PMCID: PMCPMC4825819.

18. Ruijtenberg S, van den Heuvel S. G1/S Inhibitors and the SWI/SNF Complex Control Cell-Cycle Exit during Muscle Differentiation. Cell. 2015;162(2):300–13. doi: 10.1016/j.cell.2015.06.013. PubMed PMID: 26144318.

19. Andrade-Zapata I, Baonza A. The bHLH factors extramacrochaetae and daughterless control cell cycle in Drosophila imaginal discs through the transcriptional regulation of the Cdc25 phosphatase string. PLoS Genet. 2014;10(3):e1004233. doi: 10.1371/journal.pgen.1004233. PubMed PMID: 24651265; PubMed Central PMCID: PMCPMC3961188.

20. Djiane A, Krejci A, Bernard F, Fexova S, Millen K, Bray SJ. Dissecting the mechanisms of Notch induced hyperplasia. EMBO J. 2013;32(1):60–71. doi: 10.1038/emboj.2012.326. PubMed PMID: 23232763; PubMed Central PMCID: PMCPMC3545308.

21. Jones L, Richardson H, Saint R. Tissue-specific regulation of cyclin E transcription during Drosophila melanogaster embryogenesis. Development. 2000;127(21):4619–30. PubMed PMID: 11023865.

22. Lehman DA, Patterson B, Johnston LA, Balzer T, Britton JS, Saint R, et al. Cis-regulatory elements of the mitotic regulator, string/Cdc25. Development. 1999;126(9):1793–803. PubMed PMID: 10101114.

23. Lopes CS, Casares F. Eye selector logic for a coordinated cell cycle exit. PLoS Genet. 2015;11(2):e1004981. doi: 10.1371/journal.pgen.1004981. PubMed PMID: 25695251; PubMed Central PMCID: PMCPMC4335009.

24. Bradley-Gill MR, Kim M, Feingold D, Yergeau C, Houde J, Moon NS. Alternate transcripts of the Drosophila “activator” E2F are necessary for maintenance of cell cycle exit during development. Dev Biol. 2016;411(2):195–206. doi: 10.1016/j.ydbio.2016.02.004. PubMed PMID: 26859702.

25. Dimova DK, Stevaux O, Frolov MV, Dyson NJ. Cell cycle-dependent and cell cycle-independent control of transcription by the Drosophila E2F/RB pathway. Genes Dev. 2003;17(18):2308–20. PubMed PMID: 12975318.

26. Neufeld TP, de la Cruz AF, Johnston LA, Edgar BA. Coordination of growth and cell division in the Drosophila wing. Cell. 1998;93(7):1183–93. PubMed PMID: 9657151.

27. Reis T, Edgar BA. Negative regulation of dE2F1 by cyclin-dependent kinases controls cell cycle timing. Cell. 2004;117(2):253–64. PubMed PMID: 15084262.

28. Hay BA, Wolff T, Rubin GM. Expression of baculovirus P35 prevents cell death in Drosophila. Development. 1994;120(8):2121–9. Epub 1994/08/01. PubMed PMID: 7925015.

29. Pulianmackal AJ, Kanakousaki K, Flegel K, Grushko OG, Gourley E, Rozich E, et al. Misregulation of Nucleoporins 98 and 96 leads to defects in protein synthesis that promote hallmarks of tumorigenesis. Dis Model Mech. 2022;15(3). Epub 2022/02/03. doi: 10.1242/dmm.049234. PubMed PMID: 35107131; PubMed Central PMCID: PMCPMC8938402.

30. Thacker SA, Bonnette PC, Duronio RJ. The contribution of E2F-regulated transcription to Drosophila PCNA gene function. Curr Biol. 2003;13(1):53–8. PubMed PMID: 12526745.

31. Ahringer J. NuRD and SIN3 histone deacetylase complexes in development. Trends Genet. 2000;16(8):351–6. doi: 10.1016/s0168-9525(00)02066-7. PubMed PMID: 10904264.

32. Aughey GN, Forsberg E, Grimes K, Zhang S, Southall TD. NuRD-independent Mi-2 activity represses ectopic gene expression during neuronal maturation. EMBO Rep. 2023;24(4):e55362. Epub 20230201. doi: 10.15252/embr.202255362. PubMed PMID: 36722816; PubMed Central PMCID: PMCPMC10074086.

33. Kunert N, Wagner E, Murawska M, Klinker H, Kremmer E, Brehm A. dMec: a novel Mi-2 chromatin remodelling complex involved in transcriptional repression. EMBO J. 2009;28(5):533–44. Epub 20090122. doi: 10.1038/emboj.2009.3. PubMed PMID: 19165147; PubMed Central PMCID: PMCPMC2657585.

34. Kim J, Lu C, Srinivasan S, Awe S, Brehm A, Fuller MT. Blocking promiscuous activation at cryptic promoters directs cell type-specific gene expression. Science. 2017;356(6339):717–21. doi: 10.1126/science.aal3096. PubMed PMID: 28522526; PubMed Central PMCID: PMCPMC5572561.

35. Kreher J, Kovac K, Bouazoune K, Macinkovic I, Ernst AL, Engelen E, et al. EcR recruits dMi-2 and increases efficiency of dMi-2-mediated remodelling to constrain transcription of hormone-regulated genes. Nat Commun. 2017;8:14806. doi: 10.1038/ncomms14806. PubMed PMID: 28378812; PubMed Central PMCID: PMCPMC5382322.

36. Murawska M, Hassler M, Renkawitz-Pohl R, Ladurner A, Brehm A. Stress-induced PARP activation mediates recruitment of Drosophila Mi-2 to promote heat shock gene expression. PLoS Genet. 2011;7(7):e1002206. Epub 20110728. doi: 10.1371/journal.pgen.1002206. PubMed PMID: 21829383; PubMed Central PMCID: PMCPMC3145624.

37. Fasulo B, Deuring R, Murawska M, Gause M, Dorighi KM, Schaaf CA, et al. The Drosophila MI-2 chromatin-remodeling factor regulates higher-order chromatin structure and cohesin dynamics in vivo. PLoS Genet. 2012;8(8):e1002878. doi: 10.1371/journal.pgen.1002878. PubMed PMID: 22912596; PubMed Central PMCID: PMCPMC3415455.

38. Polo SE, Kaidi A, Baskcomb L, Galanty Y, Jackson SP. Regulation of DNA-damage responses and cell-cycle progression by the chromatin remodelling factor CHD4. EMBO J. 2010;29(18):3130–9. Epub 20100806. doi: 10.1038/emboj.2010.188. PubMed PMID: 20693977; PubMed Central PMCID: PMCPMC2944064.

39. Unhavaithaya Y, Shin TH, Miliaras N, Lee J, Oyama T, Mello CC. MEP-1 and a homolog of the NURD complex component Mi-2 act together to maintain germline-soma distinctions in C. elegans. Cell. 2002;111(7):991–1002. doi: 10.1016/s0092-8674(02)01202-3. PubMed PMID: 12507426.

40. Moses K, Rubin GM. Glass encodes a site-specific DNA-binding protein that is regulated in response to positional signals in the developing Drosophila eye. Genes Dev. 1991;5(4):583–93. doi: 10.1101/gad.5.4.583. PubMed PMID: 2010085.

41. Moses K, Ellis MC, Rubin GM. The glass gene encodes a zinc-finger protein required by Drosophila photoreceptor cells. Nature. 1989;340(6234):531–6. doi: 10.1038/340531a0. PubMed PMID: 2770860.

42. Tomlinson A, Kimmel BE, Rubin GM. rough, a Drosophila homeobox gene required in photoreceptors R2 and R5 for inductive interactions in the developing eye. Cell. 1988;55(5):771–84. doi: 10.1016/0092-8674(88)90133-x. PubMed PMID: 2903798.

43. Mishra M, Oke A, Lebel C, McDonald EC, Plummer Z, Cook TA, et al. Pph13 and orthodenticle define a dual regulatory pathway for photoreceptor cell morphogenesis and function. Development. 2010;137(17):2895–904. Epub 20100728. doi: 10.1242/dev.051722. PubMed PMID: 20667913; PubMed Central PMCID: PMCPMC2938920.

44. Zhang S, Wu S, Yao R, Wei X, Ohlstein B, Guo Z. Eclosion muscles secrete ecdysteroids to initiate asymmetric intestinal stem cell division in Drosophila. Dev Cell. 2024;59(1):125–40 e12. Epub 20231213. doi: 10.1016/j.devcel.2023.11.016. PubMed PMID: 38096823; PubMed Central PMCID: PMCPMC12927586.

45. Mirth C. Ecdysteroid control of metamorphosis in the differentiating adult leg structures of Drosophila melanogaster. Dev Biol. 2005;278(1):163–74. doi: 10.1016/j.ydbio.2004.10.026. PubMed PMID: 15649469.

46. Ashburner M, Chihara C, Meltzer P, Richards G. Temporal control of puffing activity in polytene chromosomes. Cold Spring Harb Symp Quant Biol. 1974;38:655–62. doi: 10.1101/sqb.1974.038.01.070. PubMed PMID: 4208797.

47. Brennan CA, Li TR, Bender M, Hsiung F, Moses K. Broad-complex, but not ecdysone receptor, is required for progression of the morphogenetic furrow in the Drosophila eye. Development. 2001;128(1):1–11. doi: 10.1242/dev.128.1.1. PubMed PMID: 11092806.

48. Wang H, Morrison CA, Ghosh N, Tea JS, Call GB, Treisman JE. The Blimp-1 transcription factor acts in non-neuronal cells to regulate terminal differentiation of the Drosophila eye. Development. 2022;149(7). Epub 20220331. doi: 10.1242/dev.200217. PubMed PMID: 35297965; PubMed Central PMCID: PMCPMC8995086.

49. Stoiber M, Celniker S, Cherbas L, Brown B, Cherbas P. Diverse Hormone Response Networks in 41 Independent Drosophila Cell Lines. G3 (Bethesda). 2016;6(3):683–94. Epub 20160115. doi: 10.1534/g3.115.023366. PubMed PMID: 26772746; PubMed Central PMCID: PMCPMC4777130.

50. Uyehara CM, Leatham-Jensen M, McKay DJ. Opportunistic binding of EcR to open chromatin drives tissue-specific developmental responses. Proc Natl Acad Sci U S A. 2022;119(40):e2208935119. Epub 20220926. doi: 10.1073/pnas.2208935119. PubMed PMID: 36161884; PubMed Central PMCID: PMCPMC9546573.

51. Mou X, Duncan DM, Baehrecke EH, Duncan I. Control of target gene specificity during metamorphosis by the steroid response gene E93. Proc Natl Acad Sci U S A. 2012;109(8):2949–54. Epub 20120202. doi: 10.1073/pnas.1117559109. PubMed PMID: 22308414; PubMed Central PMCID: PMCPMC3286987.

52. Lam G, Nam HJ, Velentzas PD, Baehrecke EH, Thummel CS. Drosophila E93 promotes adult development and suppresses larval responses to ecdysone during metamorphosis. Dev Biol. 2022;481:104–15. Epub 20211011. doi: 10.1016/j.ydbio.2021.10.001. PubMed PMID: 34648816; PubMed Central PMCID: PMCPMC8665130.

53. Niederhuber MJ, Leatham-Jensen M, McKay DJ. The SWI/SNF nucleosome remodeler constrains enhancer activity during Drosophila wing development. Genetics. 2024;226(2). doi: 10.1093/genetics/iyad196. PubMed PMID: 37949841; PubMed Central PMCID: PMCPMC10847718.

54. Palijan A, Fernandes I, Verway M, Kourelis M, Bastien Y, Tavera-Mendoza LE, et al. Ligand-dependent corepressor LCoR is an attenuator of progesterone-regulated gene expression. J Biol Chem. 2009;284(44):30275–87. Epub 20090910. doi: 10.1074/jbc.M109.051201. PubMed PMID: 19744932; PubMed Central PMCID: PMCPMC2781583.

55. Ma Y, Kanakousaki K, Buttitta L. How the cell cycle impacts chromatin architecture and influences cell fate. Front Genet. 2015;6:19. doi: 10.3389/fgene.2015.00019. PubMed PMID: 25691891; PubMed Central PMCID: PMCPMC4315090.

56. Ma Y, Buttitta L. Chromatin organization changes during the establishment and maintenance of the postmitotic state. Epigenetics Chromatin. 2017;10(1):53. Epub 2017/11/12. doi: 10.1186/s13072-017-0159-8. PubMed PMID: 29126440; PubMed Central PMCID: PMCPMC5681785.

57. Brehm A, Langst G, Kehle J, Clapier CR, Imhof A, Eberharter A, et al. dMi-2 and ISWI chromatin remodelling factors have distinct nucleosome binding and mobilization properties. EMBO J. 2000;19(16):4332–41. doi: 10.1093/emboj/19.16.4332. PubMed PMID: 10944116; PubMed Central PMCID: PMCPMC302042.

58. Bouazoune K, Mitterweger A, Langst G, Imhof A, Akhtar A, Becker PB, et al. The dMi-2 chromodomains are DNA binding modules important for ATP-dependent nucleosome mobilization. EMBO J. 2002;21(10):2430–40. doi: 10.1093/emboj/21.10.2430. PubMed PMID: 12006495; PubMed Central PMCID: PMCPMC125999.

59. Xue Y, Wong J, Moreno GT, Young MK, Cote J, Wang W. NURD, a novel complex with both ATP-dependent chromatin-remodeling and histone deacetylase activities. Mol Cell. 1998;2(6):851–61. doi: 10.1016/s1097-2765(00)80299-3. PubMed PMID: 9885572.

60. Baehrecke EH, Thummel CS. The Drosophila E93 gene from the 93F early puff displays stage- and tissue-specific regulation by 20-hydroxyecdysone. Dev Biol. 1995;171(1):85–97. doi: 10.1006/dbio.1995.1262. PubMed PMID: 7556910.

61. Nystrom SL, Niederhuber MJ, McKay DJ. Expression of E93 provides an instructive cue to control dynamic enhancer activity and chromatin accessibility during development. Development. 2020;147(6). Epub 20200316. doi: 10.1242/dev.181909. PubMed PMID: 32094114; PubMed Central PMCID: PMCPMC7097197.

62. Martin BJE, Ablondi EF, Goglia C, Mimoso CA, Espinel-Cabrera PR, Adelman K. Global identification of SWI/SNF targets reveals compensation by EP400. Cell. 2023;186(24):5290–307 e26. Epub 20231102. doi: 10.1016/j.cell.2023.10.006. PubMed PMID: 37922899; PubMed Central PMCID: PMCPMC11307202.

63. Flegel K, Grushko O, Bolin K, Griggs E, Buttitta L. Roles for the Histone Modifying and Exchange Complex NuA4 in Cell Cycle Progression in Drosophila melanogaster. Genetics. 2016;203(3):1265–81. doi: 10.1534/genetics.116.188581. PubMed PMID: 27184390; PubMed Central PMCID: PMCPMC4937485.

64. Buchert EM, Fogarty EA, Uyehara CM, McKay DJ, Buttitta LA. A tissue dissociation method for ATAC-seq and CUT&RUN in Drosophila pupal tissues. Fly (Austin). 2023;17(1):2209481. doi: 10.1080/19336934.2023.2209481. PubMed PMID: 37211836; PubMed Central PMCID: PMCPMC10208176.

65. Robinson MD, McCarthy DJ, Smyth GK. edgeR: a Bioconductor package for differential expression analysis of digital gene expression data. Bioinformatics. 2010;26(1):139–40. Epub 20091111. doi: 10.1093/bioinformatics/btp616. PubMed PMID: 19910308; PubMed Central PMCID: PMCPMC2796818.

66. Dobin A, Davis CA, Schlesinger F, Drenkow J, Zaleski C, Jha S, et al. STAR: ultrafast universal RNA-seq aligner. Bioinformatics. 2013;29(1):15–21. Epub 2012/10/30. doi: 10.1093/bioinformatics/bts635. PubMed PMID: 23104886; PubMed Central PMCID: PMCPMC3530905.

67. Ramirez F, Ryan DP, Gruning B, Bhardwaj V, Kilpert F, Richter AS, et al. deepTools2: a next generation web server for deep-sequencing data analysis. Nucleic Acids Res. 2016;44(W1):W160–5. Epub 2016/04/16. doi: 10.1093/nar/gkw257. PubMed PMID: 27079975; PubMed Central PMCID: PMCPMC4987876.

68. Quinlan AR, Hall IM. BEDTools: a flexible suite of utilities for comparing genomic features. Bioinformatics. 2010;26(6):841–2. Epub 20100128. doi: 10.1093/bioinformatics/btq033. PubMed PMID: 20110278; PubMed Central PMCID: PMCPMC2832824.

69. Gel B, Diez-Villanueva A, Serra E, Buschbeck M, Peinado MA, Malinverni R. regioneR: an R/Bioconductor package for the association analysis of genomic regions based on permutation tests. Bioinformatics. 2016;32(2):289–91. Epub 20150930. doi: 10.1093/bioinformatics/btv562. PubMed PMID: 26424858; PubMed Central PMCID: PMCPMC4708104.

